# *In vivo* super-resolution track-density imaging for thalamic nuclei identification

**DOI:** 10.1101/2021.01.03.425122

**Authors:** Gianpaolo Antonio Basile, Salvatore Bertino, Alessia Bramanti, Giuseppe Pio Anastasi, Demetrio Milardi, Alberto Cacciola

## Abstract

The development of novel techniques for the *in vivo*, non-invasive visualization and identification of thalamic nuclei has represented a major challenge for human neuroimaging research in the last decades. Thalamic nuclei have important implications in various key aspects of brain physiology and many of them show selective alterations in various neurologic and psychiatric disorders. In addition, both surgical stimulation and ablation of specific thalamic nuclei have been proven to be useful for the treatment of different neuropsychiatric diseases. The present work aimed at describing a novel protocol for histologically-guided delineation of thalamic nuclei based on short-tracks track-density imaging (stTDI), which is an advanced imaging technique that exploits high angular resolution diffusion tractography to obtain super-resolved white matter maps with high anatomical information. We tested this protocol on i) six healthy individual 3T MRI scans from the Human Connectome Project database, and on ii) a group population template reconstructed by averaging 100 unrelated healthy subjects scans from the same repository. We demonstrated that this approach can identify up to 13 distinct thalamic nuclei bilaterally with very high reliability (intraclass correlation coefficient: 0.996, 95% CI: 0.993-0.998; total accumulated overlap: 0.43) and that both subject-based and group-level thalamic parcellation show a fair share of similarity to a recent standard-space histological thalamic atlas. Finally, we showed that stTDI-derived thalamic maps can be successfully employed to study thalamic structural and functional connectivity, and may have potential implications both for basic and translational research, as well as for pre-surgical planning purposes.

## Introduction

Thalamic nuclei show distinct and peculiar cyto-, myelo- and recepto-architectonic features, connectivity profiles and functional implications (Hirai and Jones 1989; Stepniewska et al. 1994; Percheron et al. 1996; Morel et al. 1997; Kultas-Ilinsky et al. 2003; Percheron 2003; Fang et al. 2006). Although the nomenclature of thalamic structures is still a matter of lively debate (Krack et al. 2002; Mai and Forutan 2012; Ilinsky et al. 2018; Mai and Majtanik 2019), thalamic nuclei can be traditionally subdivided into i) midline and intralaminar group, involved in limbic circuitry and arousal (Van der Werf et al. 2002; Vertes et al. 2015); ii) an anterior group, that is part of the hippocampal circuitry and has a role in episodic memory (Child and Benarroch 2013); iii) a medial compartment, whose most prominent nucleus, the mediodorsal (MD) nucleus, plays a major role in cognitive functions and in the ventral striatal circuitry (Groenewegen et al. 1993; Mitchell and Dalrymple-Alford 2005; Watanabe and Funahashi 2012); iv) a ventral compartment, which includes the so-called motor thalamus (Percheron et al. 1996; Ilinsky et al. 2018), involved in the basal ganglia/cerebellar circuitry (Alexander et al. 1986; Milardi et al. 2019) as well as the ventral posterior nucleus, a main relay for the somatic sensation pathway (Padberg et al. 2009); v) the pulvinar, located posteriorly, which represents the largest thalamic nucleus and has a prominent associative role (Benarroch 2015); finally, vi) the large metathalamic nuclei (lateral and medial geniculate nuclei), main relay stations in the visual and auditory pathways (Winer 1984; Derrington 2001).

Considering the crucial role played by thalamic nuclei in several higher-order brain functions, it is not surprising that most of them show also selective alteration in different brain diseases, ranging from Parkinson’s disease (PD) and tremor syndromes (Henderson 2000; Milosevic et al. 2018) to schizophrenia (Andreasen 1997; Pergola et al. 2015), Alzheimer’s disease (AD) (Braak and Braak 1991) and multiple sclerosis (MS) (Planche et al. 2020).

In addition, thalamic nuclei have been increasingly used as target in functional neurosurgery for the treatment of PD and essential tremor (Cury et al. 2017), epilepsy (Fisher et al. 2010; Son et al. 2016), Gilles de la Tourette’s syndrome (GTS) (Ackermans et al. 2011) and chronic pain (Young et al. 1994). However, thalamic nuclei show very poor contrast on conventional T1- or T2-weighted structural MRI scans (Deoni et al. 2005), preventing their direct reconstruction and identification in the human brain *in vivo.* A well-consolidated approach for the identification of thalamic nuclei is based on the realization of histological atlases from manual segmentation of stacks of serial histological microphotographs. The resulting maps can be then digitalized, reconstructed in three dimensions, registered to a standard MRI template and then fitted to subject’s individual anatomy using nonlinear transformations (Chakravarty et al. 2006; Krauth et al. 2010; Sadikot et al. 2011; Iglesias et al. 2018a; Ilinsky et al. 2018). Such approach brings forward the advantage of very high anatomical resolution and specificity, while not being able to account for inter-subject anatomical variability, as histological images are often obtained from a few anatomical specimens. On the other hand, different advanced multimodal imaging methods based on structural MRI (Deoni et al. 2005; Sudhyadhom et al. 2009; Traynor et al. 2011; Tourdias et al. 2014; Su et al. 2019; Liu et al. 2020), diffusion MRI (Kumar et al. 2015; Battistella et al. 2017; Najdenovska et al. 2018), and connectivity-based analysis of both tractography and resting-state functional MRI (fMRI) (Behrens et al. 2003; Johansen-Berg et al. 2005; Zhang et al. 2008, 2010; Ji et al. 2016; Kumar et al. 2017; Lambert et al. 2017; van Oort et al. 2018) have been developed in the last decades. These methods, while offering the possibility of non-invasive, *in vivo* and individualized thalamic mapping, often yield inconsistent results when compared with each other (Iglehart et al. 2020) and are inherently limited by the lower anatomical resolution when compared to histological imaging, thus preventing the identification of smaller nuclei and resulting, in some cases, in an oversimplification of thalamic morphology (Ilinsky et al. 2018).

Track-density imaging (TDI) is a novel imaging method based on diffusion tractography. Instead of exploiting information from streamline tracking to reconstruct specific fiber bundles, or on a whole-brain level, as a quantitative measure of neural “connectivity”, TDI maps the number of streamlines passing through a given grid element, as it is or by assigning to each streamline a user-defined weight that can be based on other imaging-derived measures (Calamante, Tournier, Smith, et al. 2012a; Calamante 2017). Combining diffusion MRI datasets with tractography, such approach allows to obtain high-quality images that are characterized by a high anatomical contrast of grey and white matter structures as well as super-resolution properties (up to 0.25 mm) (Calamante et al. 2010, 2011). However, despite its great potential for *in vivo* thalamic mapping, just one early work (Calamante et al. 2013) used whole-brain TDI obtained from a 7T MRI dataset to identify thalamic structures. Although these findings suggest the feasibility of such approach, this method has never been tested on 3T MRI data, which are more commonly available worldwide and in the clinical practice.

In the present paper, we firstly describe a novel histology-guided protocol based on optimized super-resolution (0.25mm) short-tracks TDI (stTDI) (Calamante, Tournier, Kurniawan, et al. 2012; Dai et al. 2018) for the identification of thalamic nuclei, demonstrating that such method can be successfully applied on single subject datasets to achieve an individualized and reliable thalamic parcellation at 3T field strength. Then, we test this approach on a population-level template derived from diffusion MRI data of 100 healthy subjects from the Human Connectome Project (HCP) (Van Essen et al. 2013).

Third, we compare our parcellation results with a previously available thalamic histological atlas (Ilinsky et al. 2018), showing a fair share of similarity to both population template-level and subject-level parcellations. Finally, a comprehensive, qualitative and quantitative structural and functional connectomic analysis of the identified thalamic nuclei is provided.

## Materials and methods

### Subjects

We employed minimally preprocessed structural, diffusion and resting-state functional MRI datasets of 210 healthy subjects (males=92, females=118, age range 22-36 years) from the HCP repository. Data have been acquired by the Washington University, University of Minnesota and Oxford university (WU-Minn) HCP consortium. The Washington University in St. Louis Institutional Review Board (IRB) (37) approved subject recruitment procedures, informed consent and sharing of de-identified data.

### Data acquisition and preprocessing

MRI data were acquired on a custom-made Siemens 3T “Connectome Skyra” (Siemens, Erlangen, Germany), provided with a Siemens SC72 gradient coil and maximum gradient amplitude (Gmax) of 100 mT/m (initially 70 mT/m and 84 mT/m in the pilot phase), to improve acquisitions of diffusion-weighted imaging (DWI).

High resolution T1-weighted MPRAGE images were collected using the following parameters: voxel size = 0.7 mm, TE = 2.14 ms, TR = 2,400 ms (Van Essen et al. 2012a).

DWI acquisition was carried out using a single-shot 2D spin-echo multiband Echo Planar Imaging (EPI) sequence and equally distributed over 3 shells (b-values 1000, 2000, 3000 mm/s2), 90 directions per shell, spatial isotropic resolution of 1.25 mm (Sotiropoulos et al. 2013). Data are available in a minimally preprocessed form that includes EPI susceptibility-based distortion and motion correction, as well as co-registration of structural and DWI images (Glasser et al. 2013).

Resting state functional MRI scans (rsfMRI) were acquired using gradient-echo EPI with 2.0 mm isotropic resolution (FOV = 208× 180mm, matrix = 104 × 90, 72 slices, TR = 720ms, TE = 33.1ms, FA =52°, multi-band factor = 8, 1200 frames, ~15 min/run) (Van Essen et al. 2012b; Ugurbil et al. 2013). Scans were acquired along two different sessions on different days, with each session consisting of a left-to-right and a right-to-left phase encoding acquisition (Van Essen et al. 2012a; Smith et al. 2013); in the present work we employ left-to-right and right-to-left acquisitions from a single session only (first session). Minimal preprocessing for rsfMRI included the following steps: artifact correction, motion correction, registration to standard space (MNI152), high pass temporal filtering (> 2,000 s full width at half maximum) for removal of slow drifts (Glasser et al. 2013), artifact components identification using ICA-FIX (Salimi-Khorshidi et al. 2014) and regression of artifacts and motion-related parameters (Smith et al. 2013).

### DWI post-processing and diffusion signal modeling

T1-weighted structural images underwent brain extraction, cortical and subcortical segmentation implemented by BET, FAST and FIRST FSL functions respectively (https://fsl.fmrib.ox.ac.uk/fsl) to obtain five tissues type images (5TT) which were employed to run multi-shell, multi-tissue constrained spherical deconvolution (MSMT-CSD), an optimized version of the CSD signal modeling, which estimates white matter Fiber Orientation Distribution (FOD) function from the diffusion-weighted deconvolution signal using a single fiber response function (RF) as reference (Tournier et al. 2008, 2012). MSMT-CSD improves the classical CSD approach calculating different response functions for gray matter, white matter and cerebrospinal fluid (Jeurissen et al. 2014a).

### Tractography and stTDI generation

An unbiased population-based FOD template was built using FOD datasets estimated from 100 unrelated subjects (males = 46, females = 54, age range 22-36 years); the template was optimized by iterating a symmetric, unbiased FOD registration algorithm which includes FOD reorientation using apodised delta functions (Raffelt et al. 2011, 2012; Pietsch et al. 2017) (Figure 1A). The FOD population template was used to generate a group-level stTDI.

We also randomly selected 6 subjects (3 males, 3 females) to generate individual stTDI for subject-level thalamic parcellation (HCP IDs: 100307, 100406, 101107, 101309, 101915, 103414).

Since TDI signal-to-noise ratio is known to rely mostly on the high number of streamlines selected (Calamante 2017), we restricted stTDI generation to a box ROI covering the diencephalic area: this would allow the generation of a larger number of streamlines in a reduced space, thus resulting in improved TDI contrast while saving computational time. In order to account for inter-individual anatomical variability, we opted for a manually drawn ROI based on anatomical landmarks, rather than using a predefined size and shape for each subject. To manually define our bounding box ROI, we started from an axial section at the level of the anterior commissure (z=0), and we drew a rectangle extending anteriorly to the edge of the putamen, and posteriorly to include the temporal horn of lateral ventricles, bilaterally; lateral boundaries were tangent to the most lateral edge of the putamen. We extended this ROI on each axial section until reaching the last slice in which the body of the caudate nucleus was visible, as superior boundary, and the section at the level of the optic chiasm as inferior boundary (Figure 1A).

A short-tracks tractogram was obtained by generating 50 millions of streamlines within this bounding box ROI using the following parameters: algorithm = *iFOD2* (Tournier et al. 2010), maximum track length=10 mm; minimum track length=5 mm; cutoff: 0.05; step size: 0.25 mm. From this short-tracks tractogram we obtained two super resolution stTDI maps with isotropic 0.25mm voxel size: the first with directionally-encoded color contrast (DEC-stTDI), the second with an apparent fiber density-based contrast (AFD-stTDI), i.e. by assigning to each streamline a weighting that is proportional to the mean FOD amplitude along the streamline itself (Figure 1B-1C) (Calamante, Tournier, Smith, et al. 2012a).

**Figure 1.**
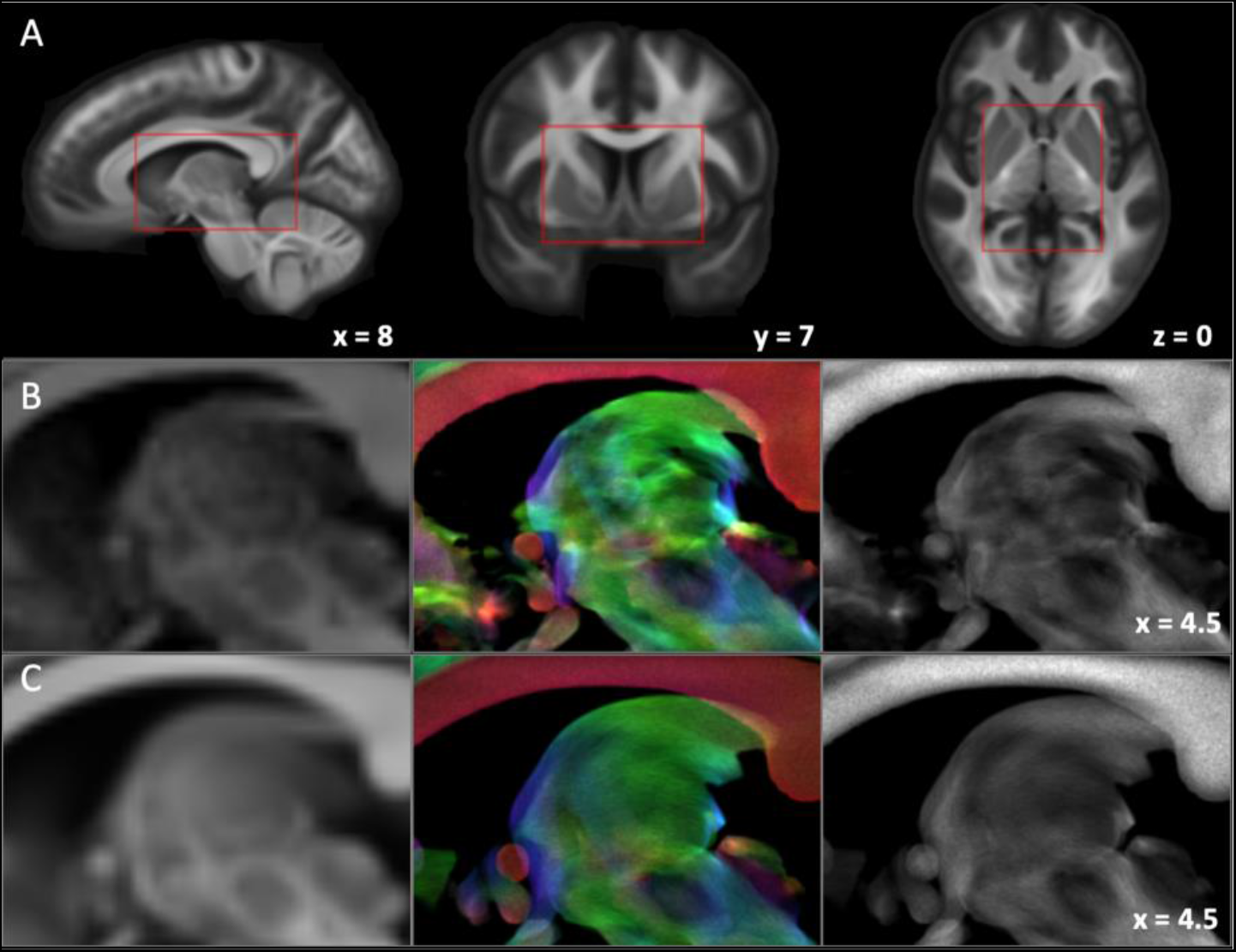
Optimized short-tracks track-density imaging (stTDI): an overview. **A)** Boundaries of the diencephalic ROI used to constrain generation of short-tracks tractogram in sagittal (left), coronal (mid) and axial (right) sections, overlaid on FOD population template. Anatomical landmarks are described in the main text. **B)** Subject-level stTDI maps. Sagittal sections of a representative subject’s FOD map (left), directionally-encoded-color stTDI (DEC-stTDI) (mid) and apparent-fiber-density stTDI (AFD-stTDI) (right). **C)** Template-level stTDI maps. Sagittal sections of FOD population template (left), DEC-stTDI (mid) and AFD-stTDI (right). Generation methods and contrast features are detailed in text.

### Histology-guided manual segmentation

Manual delineation of thalamic structures was performed on stTDI maps at the group- and individual-level by a trained neuroanatomist (D.M.). Thalamic structures were identified in sagittal slices using a histology-guided approach. Seriate 0.25 mm-thick sagittal slices of human diencephalon, Nissl-stained and with immunohistochemistry for glutamic acid decarboxilase-isoform 65 (GAD-65) were obtained from http://www.humanmotorthalamus.com/ (credited to Igor Ilinsky and Kristy Kultas-Ilinsky). Further details on histological processing are described in (Ilinsky et al. 2018).

Thalamic nuclear boundaries were delineated on each slice on DEC-stTDI maps, with AFD-stTDI as overlay (approximately around 80% opacity), starting from the midline and proceeding in medio-lateral direction, and using the histological image with the same distance to midline as reference. The obtained boundary maps were then filled to obtain 3D volumes, that were checked and further refined in axial and coronal planes using the corresponding axial and coronal histological maps (available on http://www.humanmotorthalamus.com/) as reference. Finally, volumes were median filtered (5×5×5 voxel neighborhood), eroded (number of iterations: 1) to avoid partial overlap along nuclear boundaries, binarized and resized to 0.5 mm^3^ isotropic voxel size.

### Similarity analysis

Volumes (expressed in mm^3^) have been calculated for each thalamic map derived both from group FOD template and from single-subject stTDI maps. To evaluate intra-rater reliability of volumetric estimates of thalamic nuclei derived from both subject-level and group-level parcellation, we calculated an intraclass correlation coefficient (ICC), measuring absolute agreement under a two-way random ANOVA model. All the analyses were conducted in SPSS (Version 27.0. Armonk, NY: IBM Corporation).

Thalamic nuclear maps were coregistered to standard MNI ICBM 2009b nonlinear asymmetric brain template (Fonov et al. 2009) employing ANT’s symmetric diffeomorphic nonlinear image registration tool (SyN) (Avants et al. 2008). We coregistered the FOD template using a generic affine transformation by concatenating center-of mass alignment, rigid, similarity, and fully affine transformations (similarity metric: mutual information), followed by a non-linear transformation (symmetric diffeomorphic normalization transformation model with regular sampling, similarity metric: mutual information, gradient step size: 0.2, four multi-resolution levels, smoothing sigmas: 3, 2, 1, 0 voxels [fixed image space], shrink factors: 8, 4, 2, 1 voxels [fixed image space], data winsorization [quantiles: 0.005, 0.995], convergence criterion: slope of the normalized energy profile over the past 10 iterations <10^-6^) that were concatenated into a single composite warp field. For single subject maps, we registered individual FOD maps to FOD template using ANTs SyN (same parameters as above, except for similarity metric of the non-linear transformations: cross correlation) and then we concatenated FOD to FOD template registration with FOD template to standard template registration. We used the composite warps obtained from these registration steps to coregister both group-level and subject-level thalamic maps to the standard MNI ICBM 2009b nonlinear asymmetric brain template.

Similarity measures were calculated from thalamic maps in standard space. In particular, we evaluated similarity between the group-level thalamic nuclear maps obtained from FOD population template and the Nifti version of Ilinsky’s histological human thalamic atlas featured in Lead-DBS toolbox by using Dice similarity coefficient:

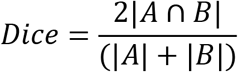

where A and B are, respectively, the number of voxels of binary thalamic maps. Similarly, we calculated Dice coefficients for each subject-level thalamic map and the histology-derived thalamic nuclei thus also obtaining an average Dice value for subject-level parcellation (Diceavg). Furthermore, we evaluated inter-subject consistency of manual thalamic segmentation (group-level and subject-level) by calculating the overlap-by-level (OBL), a measure of the overlap for each cluster across all datasets, and the total accumulated overlap (TAO), a measure of the overall, groupwise overlap across all datasets for the manual segmentation pipeline.

For a group of *m* pairs of images, where *m* represents all the possible pairwise combinations between images of the same structure, OBL is defined as:

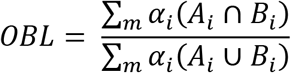

where *i* is the cluster label and α is a weighting coefficient. We defined α as the inverse of the mean of the absolute value of A and B, to avoid overestimation of larger clusters:

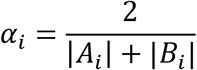

The pairwise combinations were computed including both subject-level and group-level template based thalamic parcellation, resulting then in 21 pairs. For the same group, TAO is defined as:

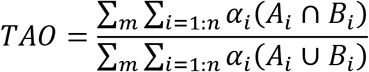

where *n* is the total number of parcels obtained from the manual segmentation (n = 13) (Traynor et al. 2010; da Silva et al. 2017).

### Structural connectivity analysis

For each thalamic nucleus obtained from manual segmentation of the template-level stTDI maps, we estimated the structural connectivity profiles of each subject (n=210) using probabilistic tractography. Population-level thalamic outlines obtained by manual segmentation were registered to each subject’s native space using ANTs SyN non-linear transformations. For each subject, a 10-million-streamlines tractogram was generated using the following parameters (algorithm = *iFOD2*, cutoff: 0.06, step size: 0.625 mm, maximum angle: 45°). Anatomically-constrained tractography (ACT) was implemented to select only streamlines terminating at the grey matter/white matter interface (Smith et al. 2012).

Quantitative connectivity was evaluated applying the Spherical Informed Filtering of Tractograms-v2 framework (SIFT2), in particular by using a combined targeted tracking/whole brain approach as described in the original publication (Smith et al. 2015). Briefly, for each thalamic nuclear ROI, 10,000 streamlines were generated using custom parameters (algorithm: iFOD2, cutoff = 0.05, step size = 1.25 mm, maximum angle = 30°, ACT); the resulting tractograms were concatenated to the previously obtained whole-brain tractogram; the mixed tractogram was fed into the SIFT2 pipeline, thus obtaining, for each streamline, a streamline weighting coefficient (*fs*) that is proportional to fiber cross sections; specific tractograms for each thalamic nucleus, along with their corresponding streamline weights, were then extracted from the mixed tractogram.

For qualitative assessment and visualization of structural connectivity profiles for each thalamic nucleus, we generated group connectivity maps with the following pipeline: first, for each tractogram, tract density maps were generated in all the subjects assigning to each streamline a contribution that is proportional to its streamline weighting coefficient (Calamante, Tournier, Smith, et al. 2012b); a threshold of 10% of the resulting track density was applied to individual maps to retain only streamlines with higher connectivity values; the resulting TDI maps were then binarized, normalized to FOD template and summed up to obtain tract maximum probability maps (MPMs); a 50% threshold was applied to MPMs to show only voxels overlapping in at least half of the sample.

In addition, we also estimated the connectivity profiles of each thalamic nucleus with brain targets in order to provide a quantitative characterization. Target ROIs included in the analysis were obtained from different commonly-available brain atlases: i) cortical parcellation was obtained from Automated Anatomic Atlas, version 3 (AAL3) (Rolls et al. 2020); ii) subcortical nuclear ROIs were extracted from Reinforcement Learning Atlas (Pauli et al. 2018); iii) cerebellar ROIs from the MNI version of the Spatially Unbiased Infratentorial Atlas (SUIT) (Diedrichsen 2006; Diedrichsen et al. 2011); iv) finally, white matter brainstem pathways from Tang et al. probabilistic atlas (Tang et al. 2018). All these ROIs were normalized from standard to subject space using nonlinear transformations. As regard the cerebellar parcellation, we considered in our analysis cerebellar cortex only, and we merged cerebellar cortical ROIs into bilateral anterior vermis (left and right lobules I-IV and V), bilateral posterior cerebellar hemispheres (left and right lobules from VI to IX); unilateral median posterior vermis (vermal lobules from VI to IX) and unilateral flocculonodular lobe (vermis and hemispheric lobule X) following the functional subdivision of cerebellar cortex employed in (Stoodley and Schmahmann 2018; Cacciola, Bertino, et al. 2019). Specific ROI-to-target tractograms for each thalamic nucleus were extracted from the mixed tractograms; the sum of streamline weights (F) for each tractogram was obtained and the mean across all subjects was employed as a structural connectivity estimate. Structural connectivity fingerprints were obtained after averaging left and right thalamic connectivity values for each nucleus; F values are reported as percentage of the total for each nucleus.

### Functional connectivity analysis

To obtain functional connectivity profiles for each thalamic nucleus, ICBM template-registered ROIs were resliced to 2mm^3^ isotropic voxel size and coregistered to the 2mm version of the MNI 152 brain template using a freely available transformation (credited to Andreas Horn, https://dx.doi.org/10.6084/m9.figshare.3502238.v1), in order to match the same reference space and voxel size of rsfMRI data. Minimally-preprocessed rsfMRI datasets were band-pass filtered (0.001-0.09 Hz) and underwent an additional denoising step by regressing out the global WM and CSF signal (Plachti et al. 2019). Functional connectivity was calculated for each subject by averaging each ROI’s time series and calculating bivariate Pearson’s correlation coefficient between the average time series and every other voxel in the brain; the resulting maps were Fisher’s R-to-Z-transformed. All the resting-state fMRI post-processing described above was carried out using the CONN toolbox (Whitfield-Gabrieli and Nieto-Castanon 2012).

The resulting first-level functional connectivity maps were concatenated for all subjects and for left and right ROIs and underwent random effects analysis using a one-sample t-test (10,000 permutations) within a threshold-free-cluster enhancement (TFCE) approach (FSL command *randomise)* (Smith and Nichols 2009). The output t-maps were masked with a TFCE-corrected p-value < 0.001.

Functional connectivity fingerprints were obtained by extracting, from the obtained thresholded maps, the highest t-statistic values within a set of standard-space, binarized target ROIs. In particular, we employed the same target ROIs that were used for structural connectivity analysis (see paragraph above), with the only exception of white matter brainstem pathways, that were excluded from the analysis; in addition, thalamic nuclear ROIs were also considered as targets. All target ROIs were resliced to isotropic voxel size of 2 mm^3^.

Figure 2 summarizes the entire pipeline.

**Figure 2.**
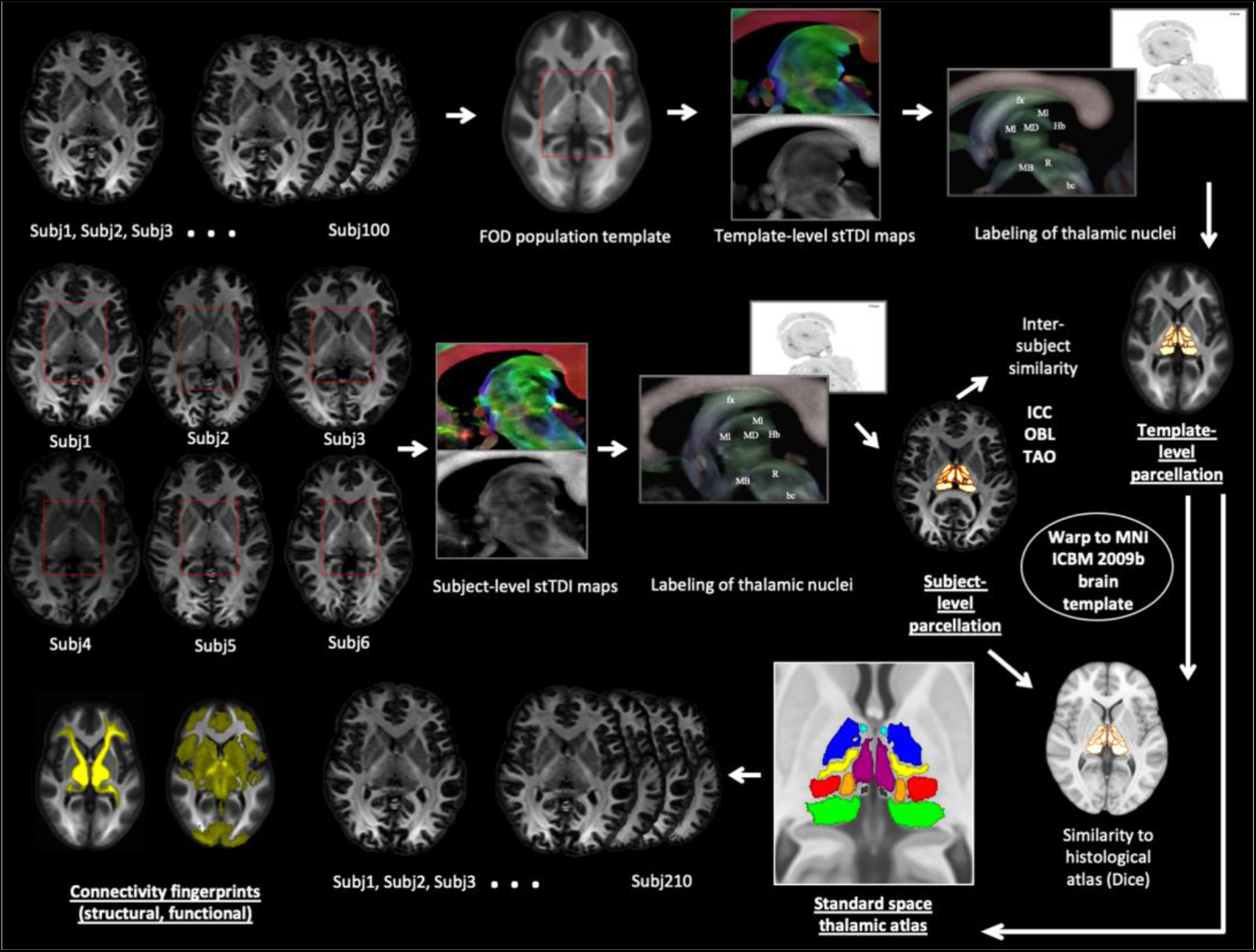
The experimental workflow in brief. The figure summarizes the experimental workflow, emphasizing template generation, the group-level and subject-level pipelines converging on the standard ICBM 2009b brain template for similarity analysis, and the following structural and functional connectivity analysis based on standard space template-level thalamic maps. All the single steps are detailed in text.

## Results

### Super-resolution and contrast properties of TDI maps

Figure 1 shows a comparison between different contrasts obtained from FOD map, DEC-stTDI and AFD-stTDI respectively both on a single representative subject (Figure 1B) and on the maps derived from group population template (Figure 1C). In the FOD map (left), intensity is proportional to the amplitude of FOD lobes and, then, white matter appears hyperintense and grey matter hypointense; as an example, in the sections showed in Figure 1A, grey matter structures such as the red nucleus, substantia nigra, superior and inferior colliculi or the thalamus itself are well contrasted from the surrounding white matter tracts, such as fornix, anterior commissure, optic tract or the mammillothalamic tract (which traverses the thalamus at the rostral level). In the FOD template (Figure 1C, left), the average between multiple scans (n=100) leads to improved contrast, which is, nevertheless, not enough to make intra-thalamic structures readily identifiable. In addition, it is worth to note that FOD maps and, subsequently, the FOD template have the same resolution of DWI scans (1.25 mm^3^). On the other hand, in stTDI maps, the super-resolution properties of stTDI allowed to achieve an isotropic voxel size of 0.25 mm^3^. In addition, the mechanism of constrained short tract generation underlying stTDI provides enhanced contrast on intra-thalamic small white matter tracts, thus allowing for the identification of hyper- and hypointense regions within the thalamus, that may likely reflect histological properties of distinct thalamic nuclei. Generally speaking, intra-laminar nuclei (that are surrounded by dense white matter tracts) are hyperintense and extra-laminar nuclei are hypointense, although there may be some exceptions to this rule. Color-coded DEC-stTDI (Figure 1B/1C, mid) provides additional information since colors are assigned to streamlines according to their directionality (red: latero-lateral; blue: superior-inferior; green: anterior-posterior). For example, the mammillothalamic tract (which is visible from anterior nucleus all the way to mammillary bodies) has a light-blue color, while anterior commissure or corpus callosum are red due to their latero-lateral orientation. This feature can be helpful to distinguish between small fiber tracts according to their different course and orientation: for example, the ansa lenticularis, running below the ventral thalamus, can be identified from the surrounding white matter for its red color. On the other hand, in AFD-stTDI contrast (Figure 1B/1C, right column), a weight is assigned to each streamline, which is proportional to the mean amplitude of the FOD lobes traversed by the streamline. In comparison to DEC-stTDI contrast, the choice of this weighting enhances the white matter vs grey matter contrast features already present in FOD maps, thus allowing a sharper distinction between hyper- and hypointense regions. These contrast properties are retained in FOD-template-based stTDI maps (Figure 1C), which also appear smoother in comparison to the “noisy” single subject maps, due to the higher signal-to-noise ratio deriving from coregistration and averaging of multiple images with same contrast features.

### Manual delineation of thalamic nuclei

Considering that contrast features alone are not always sufficient to unequivocally distinguish thalamic nuclei, comparison to matched histological sections (Figures 3-6) is crucial for identification and manual segmentation of these structures. We identified thalamic nuclei starting from 0.5mm to the midline and proceeding in a medio-lateral direction in serial sagittal slices. Table 1 summarizes the nomenclature and abbreviations adopted for labeling of histological and stTDI maps.

**Table 1.**
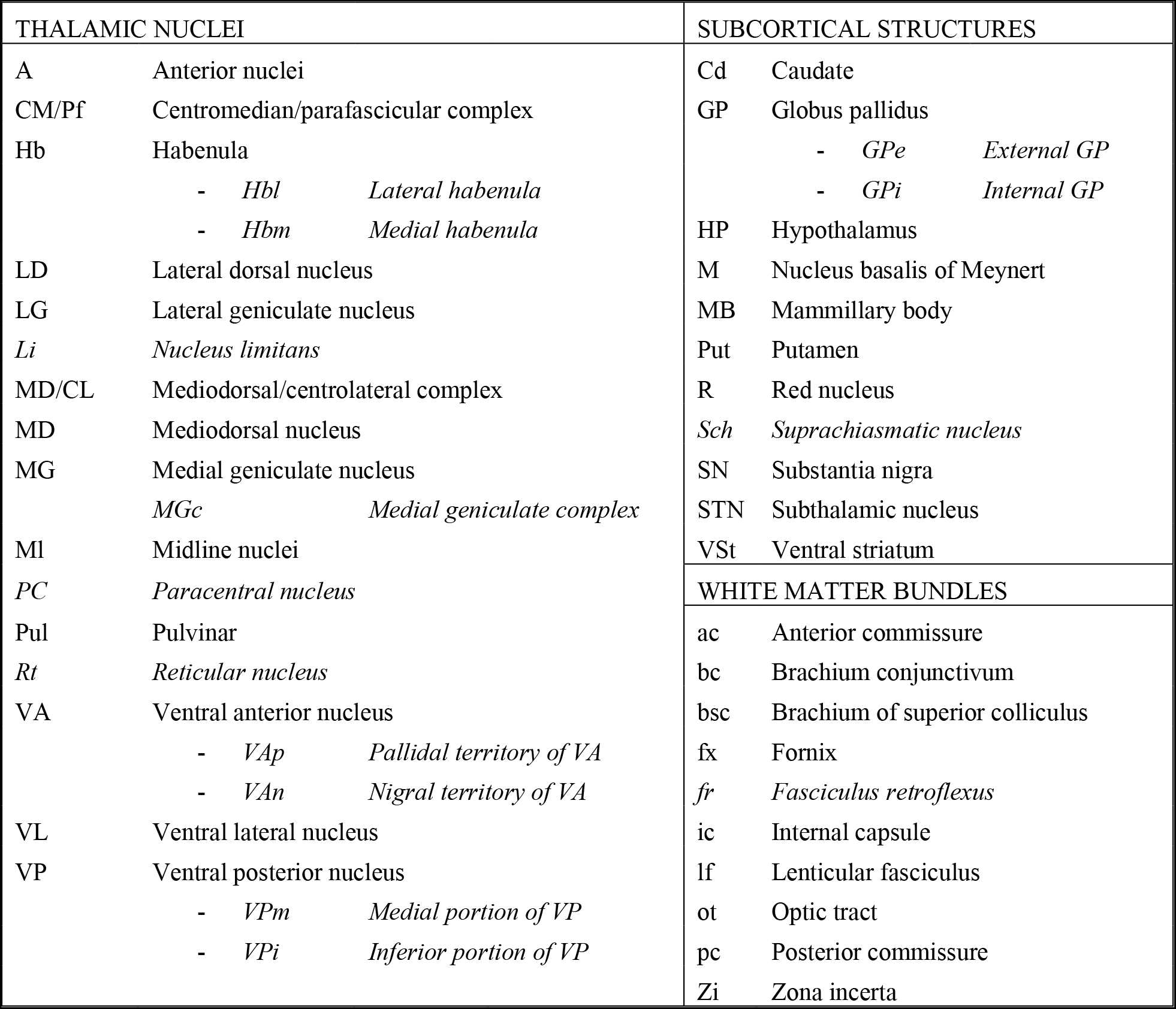
List of abbreviations used for labeling of histological and stTDI slices. Structures that were labeled on histological slices, but not on stTDI maps are reported in italics.

In the most medial sections (0.5 mm to 1.5mm) the mediodorsal (MD) nucleus is identifiable as a large, central hypointense mass surrounded by a hyperintense rim that roughly corresponds to midline (Ml) nuclei (Figure 3). Habenula (Hb) can be identified as a hyperintense region at the caudal pole of MD, which can be also visualized directly on FOD maps, and tends to a light blue color in DEC-stTDI maps. On TDI maps, we were not able to distinguish between lateral (Hbl) and medial (Hbm) habenula, as in the histological sections. In addition, in comparison to TDI and FOD maps, Hb structures appears to be displaced and collapsed downwards in an inferior and posterior position in the histological sections; this is probably caused by the loss of intrinsic CSF pressure in post-mortem preparations. Proceeding in medio-lateral sections (1.5 to 2.5mm), the anterior nucleus (A) can be identified as a hypointense region in dorsal and rostral position. It is divided from MD by a posterior-inferior hyperintense border, while the fornix (fx), that appears as a strongly hyperintense arc both in FOD and AFD-stTDI maps, marks its antero-superior border.

Posteriorly, the pulvinar (Pul) appears behind the posterior border of the MD nucleus; its posterior borders are strongly hypointense in stTDI maps, so that they can be better distinguished from surrounding CSF in FOD maps (not shown in figures).

Moving to approximately 2.5 mm from midline, the centromedian/parafascicular complex (CM/Pf) can be identified as a slightly hyperintense, oval-shaped region with a hypointense posterior tip lying just below the (hypointense) MD nucleus; a thin line of white matter (hyperintense) separates it from the red nucleus (R) on its inferior border. Due to difficulty in individuating its borders, we excluded the nucleus limitans (Li), which lies posteriorly to CM/Pf complex, from our manual parcellation protocol. Proceeding at 3.5 mm from midline, the ventral anterior nucleus (VA) becomes visible as a hypointense region lying in the anterior portion of the thalamus, just below the A nucleus, from which is divided by a thin, fuzzy hyperintense border. Notice that stTDI contrast does not allow to distinguish between nigral and pallidal afferent territories of VA nucleus (VAn and VAp respectively in the histological sections).

The lateral dorsal nucleus (LD) can be also identified as an ill-delimited hyperintense region situated behind the most caudal tip of the A nucleus, and included below the fornix superiorly (hyperintense) and the MD nucleus inferiorly (hypointense). We were not able to identify consistently the paracentral (PC) thalamic nucleus, which was then excluded from our parcellation protocol.

In slices from 3.5mm to 5.5mm the mammillothalamic tract (mt) is visible, being strongly hyperintense in FOD maps, slightly hyperintense in AFD-stTDI and with a light blue color in DEC-stTDI. The mt tract represents an important reference point since, in the most medial slices (around 3.5mm from midline), it runs through the posterior part of VA very close to its posterior border; in slices about 4.5mm from midline it can be used as a good indicator of the position of the boundary that divides the VA nucleus (hypointense) from the VL nucleus (hyperintense).

**Figure 3.**
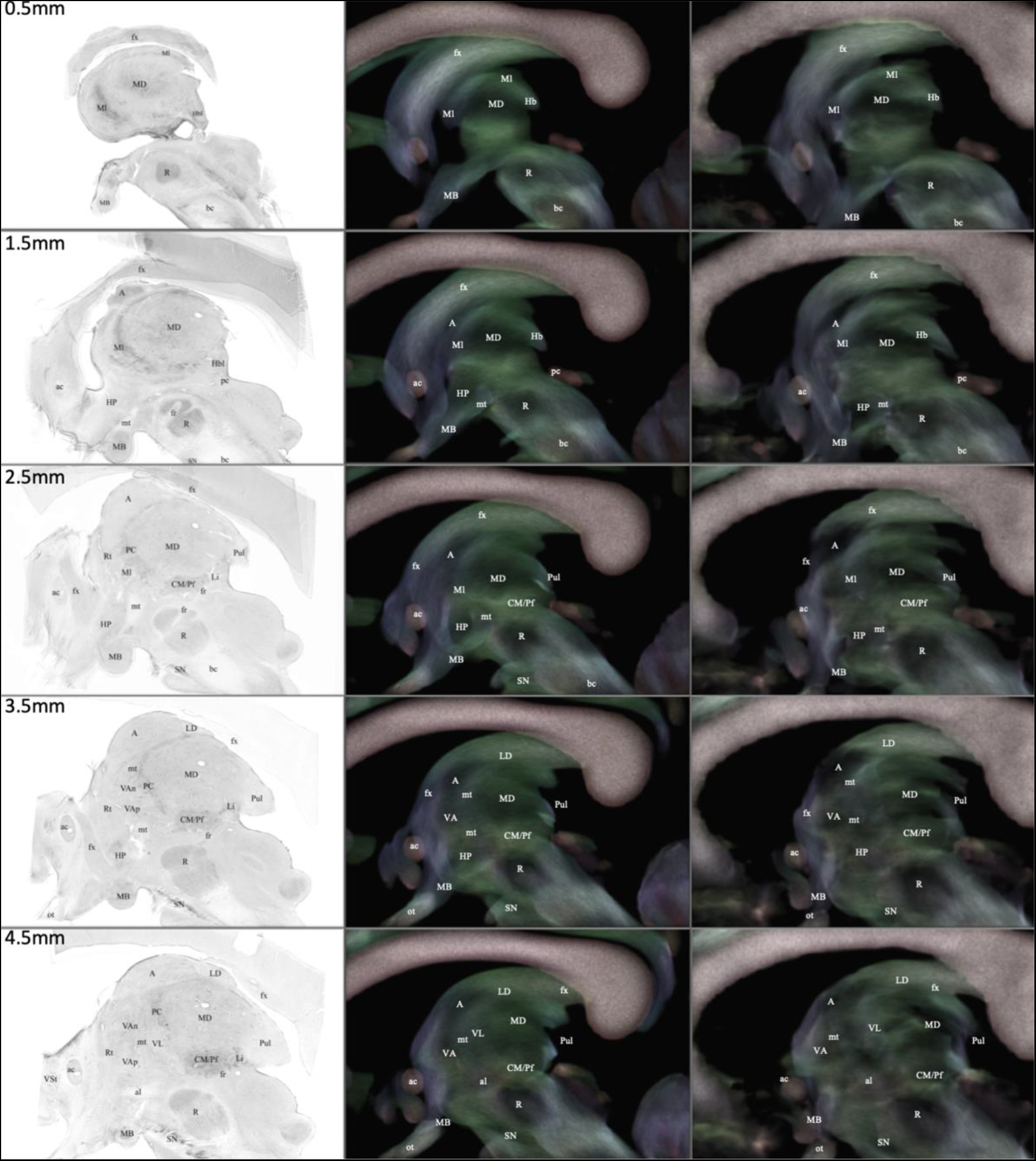
Histologically-guided labeling of thalamic structures. Slices from 0.5 to 4.5 mm from midline. Sagittal slices are derived from: histological microphotographs, Nissl/GAD-65 staining (from http://www.humanmotorthalamus.com/) (left); template-level stTDI (mid); subject-level stTDI (right). Apparent fiber density-weighted stTDI (AFD-stTDI) are overlaid on directionally-encoded-color stTDI with 80% opacity. Abbreviations are explained in Table 1. Description of structures is detailed in text.

In slices from 5.5mm to 6.5mm from midline (Figure 4), VL is increasingly visible and can be identified as a slightly hyperintense strip (green-colored in DEC-stTDI) in between the (predominantly hypointense) VA and MD nuclei. In addition, the medial portion of ventral posterior nucleus (VP) (marked as VPm in histological slices) appears as a hypointense region anterior to the CM/Pf complex, inferior to VL and posterior to VA and mt tract.

Below the VA nucleus, the subthalamic nucleus (STN) becomes visible as a hypointense region sitting on top of the substantia nigra (SN) and separated from the VA nucleus by the hyperintense ansa lenticularis (al). Around 7.5mm from midline, the A nucleus is not visible anymore. The MD nucleus becomes progressively more intense in stTDI maps, probably corresponding to the passage to its intralaminar portion, the mediodorsal/centrolateral complex (MD/CL). In slices from 7.5 to 11.5 mm, this structure is visible as an approximately triangular region situated posteriorly to VL nucleus, anteriorly to the Pul and above the CM/Pf; this region has a green to light blue color in DEC-stTDI maps. On the other hand, the CM/Pf complex (marked simply as CM in histological slices) becomes progressively more hypointense and is surrounded by a dark blue/purple hyperintense border in DEC-stTDI maps. Proceeding in medio-lateral directions, the boundaries between VA (hypointense) and VL (hyperintense) become increasingly less evident as these two nuclei tend to display similar intensities in both DEC- and AFD-stTDI maps; at this level, the histological boundary between VA and VL is slightly curved, i.e. has a crescent shape, that can be followed from the most posterior tip of STN; hence, a putative boundary between VA and VL can be approximately defined by tracing a curved vertical line tangent to the most posterior tip of the STN.

Moving to 9.5mm from midline, the medial geniculate (MG) becomes visible as a small, wedge-shaped hypointense region below the Pul, posteriorly to VP and CM/Pf and anteriorly to the brachium of superior colliculus (bsc); this latter is strongly hyperintense in AFD-stTDI maps and has a marked red color in DEC-stTDI, being composed mostly of latero-laterally oriented fibers.

**Figure 4.**
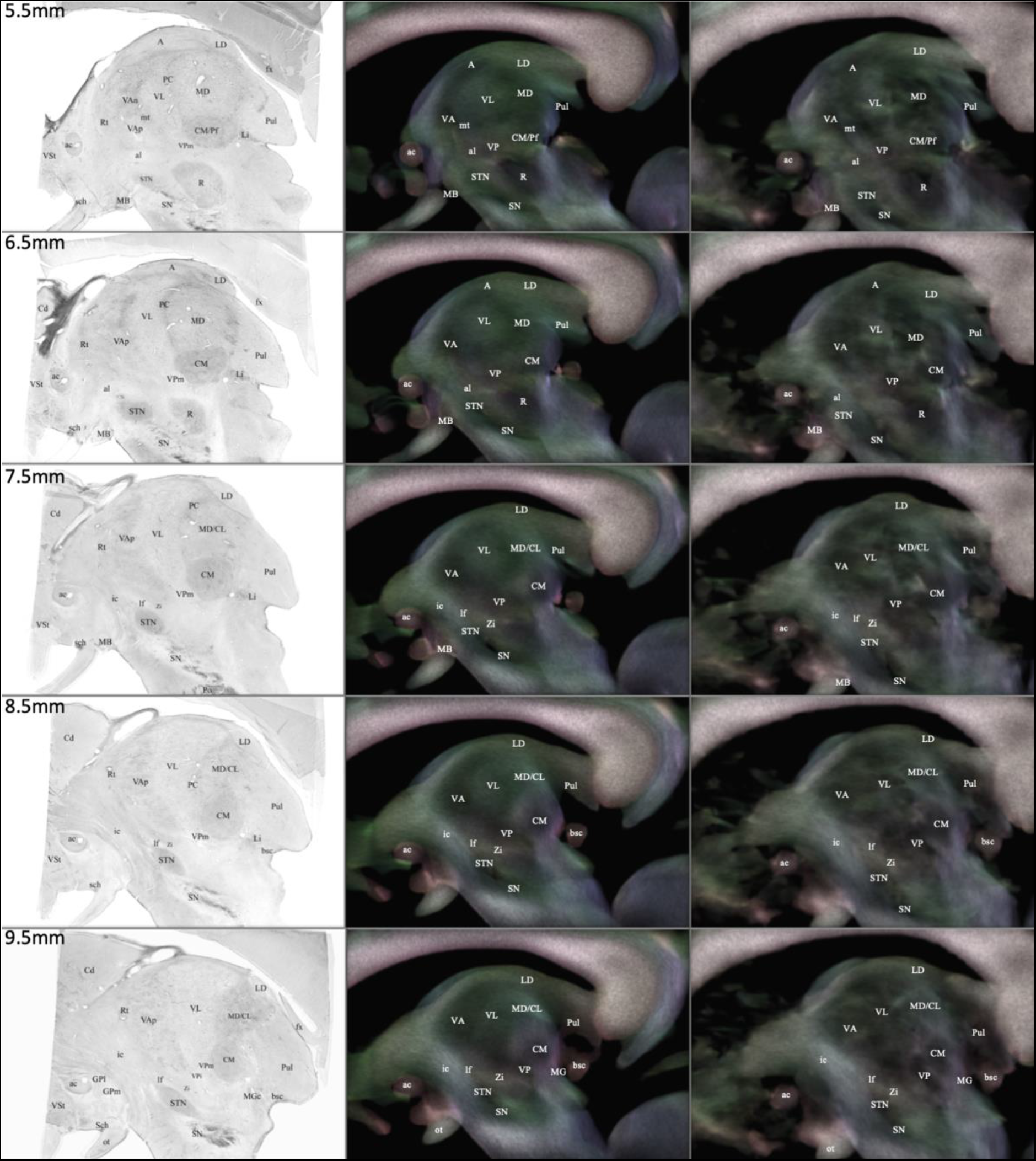
Histologically-guided labeling of thalamic structures. Slices from 5.5 to 9.5 mm from midline. Sagittal slices are derived from: histological microphotographs, Nissl/GAD-65 staining (from http://www.humanmotorthalamus.com/) (left); template-level stTDI (mid); subject-level stTDI (right). Apparent fiber density-weighted stTDI (AFD-stTDI) are overlaid on directionally-encoded-color stTDI with 80% opacity. Abbreviations are explained in Table 1. Description of structures is detailed in text.

In slices around 10.5mm from midline (Figure 5), LD nucleus is not visible anymore and CM/Pf complex becomes increasingly smaller; its boundaries tend to become gradually less distinguishable from those of VP, which is located anteriorly and has a virgule shape; at 11.5 mm from midline both CM/Pf and MD/CL are not identifiable anymore.

In slices from 11.5 to 15.5mm, five thalamic structures can be identified: VA and VL that have similar contrast properties (hypointense and with a greenish hue on DEC-stTDI maps) and are located anteriorly, just behind the internal capsule (ic); Pul, that lies posteriorly, has a drop-like shape and a reddish color on DEC-TDI; VP, that lies between VL and Pul in a more ventral position and is surrounded by slightly hyperintense, purple-colored borders in DEC-stTDI; and MG, that has a wedge-like shape, its vertex directed forward and upwards, and lies postero-inferiorly to VP and antero-inferiorly to the Pul.

**Figure 5.**
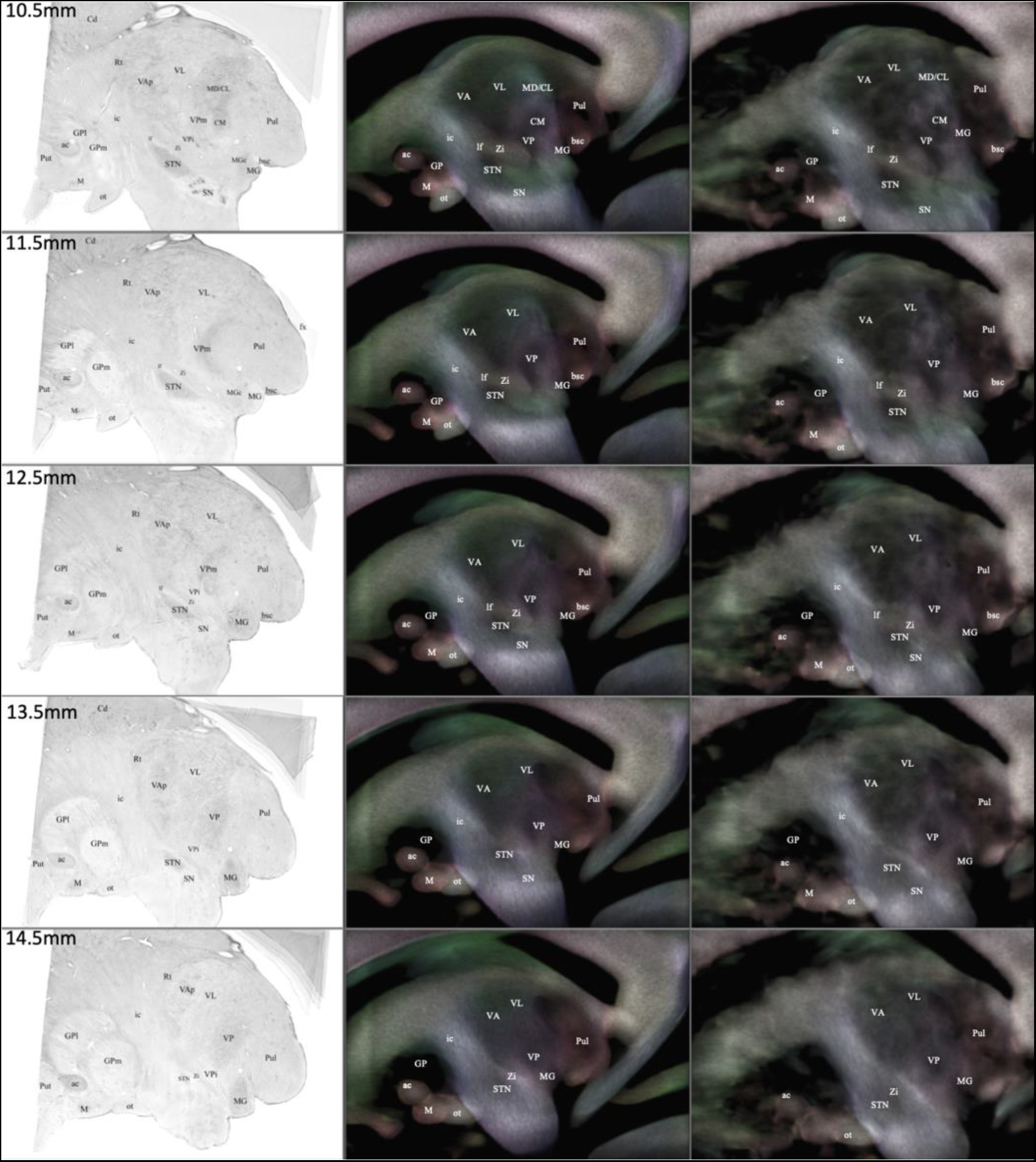
Histologically-guided labeling of thalamic structures. Slices from 10.5 to 14.5 mm from midline. Sagittal slices are derived from: histological microphotographs, Nissl/GAD-65 staining (from http://www.humanmotorthalamus.com/) (left); template-level stTDI (mid); subject-level stTDI (right). Apparent fiber density-weighted stTDI (AFD-stTDI) are overlaid on directionally-encoded-color stTDI with 80% opacity. Abbreviations are explained in Table 1. Description of structures is detailed in text.

Proceeding in the most lateral slices (Figure 6), VA and VL gradually fade into one another and disappear behind the ic, until only VP remains visible as a round-like, hypointense structure with fuzzy borders, anterior to Pul. In slices about 16.5mm from midline, MG becomes increasingly smaller until it disappears, and in the last slices is replaced by lateral geniculate nucleus (LG); this structure occupies the same position but looks considerably larger, and in the most lateral slices (around 18.5/19.5 mm from midline) it is joined by the optic tract (ot), which penetrates it at the rostral and ventral level. In the last sections (19.5mm from midline), only Pul and LG are visible and are clearly distinguished one from another; we were able to delineate Pul approximately until 20.5mm from midline, and LG until 23.5mm (not shown in figures).

**Figure 6.**
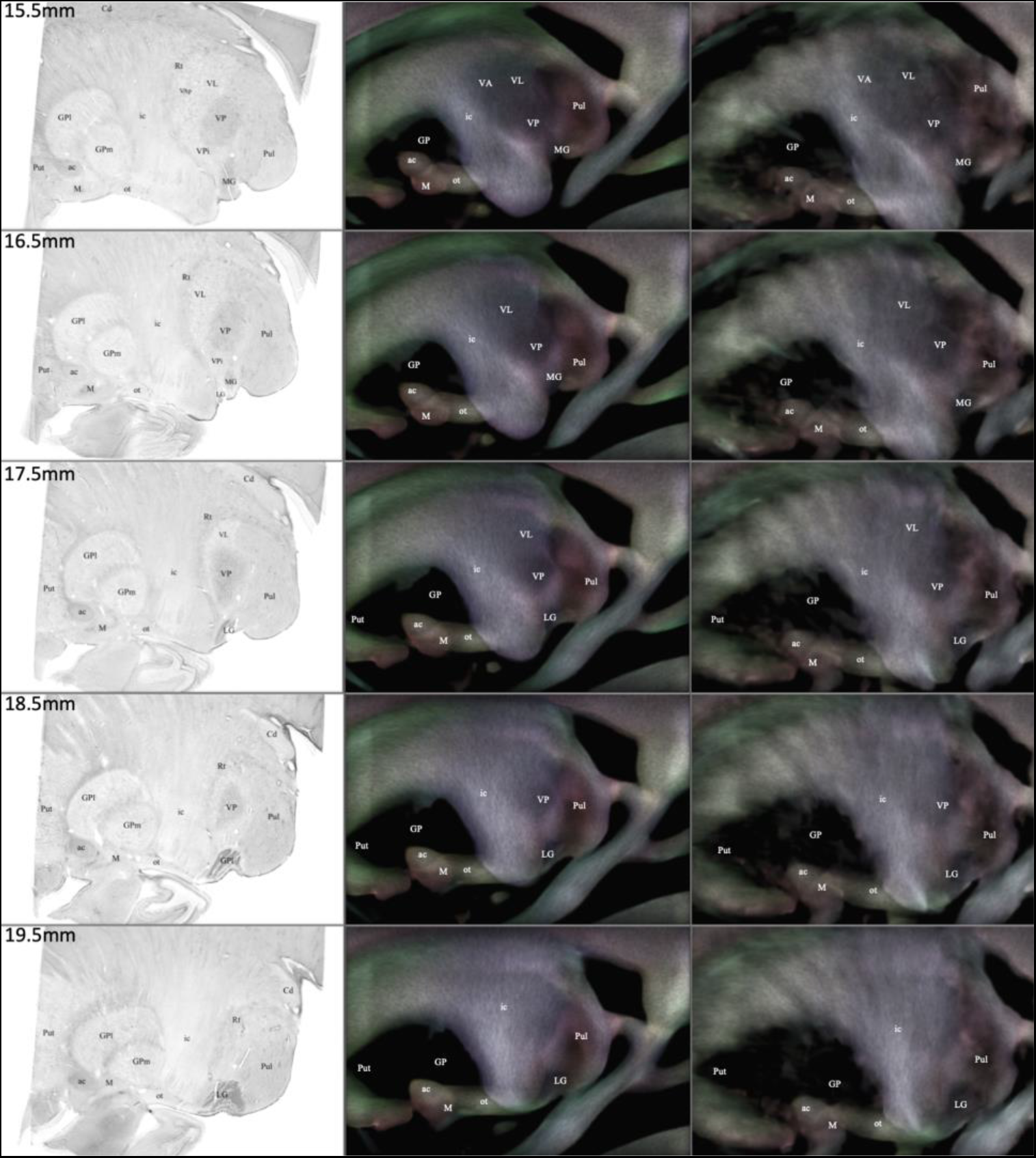
Histologically-guided labeling of thalamic structures. Slices from 15.5 to 19.5 mm from midline. Sagittal slices are derived from: histological microphotographs, Nissl/GAD-65 staining (from http://www.humanmotorthalamus.com/) (left); template-level stTDI (mid); subject-level stTDI (right). Apparent fiber density-weighted stTDI (AFD-stTDI) are overlaid on directionally-encoded-color stTDI with 80% opacity. Abbreviations are explained in Table 1. Description of structures is detailed in text.

A 3D reconstruction and 2D axial, coronal and sagittal slice representation of the template-level thalamic atlas is displayed in Figure 7.

**Figure 7.**
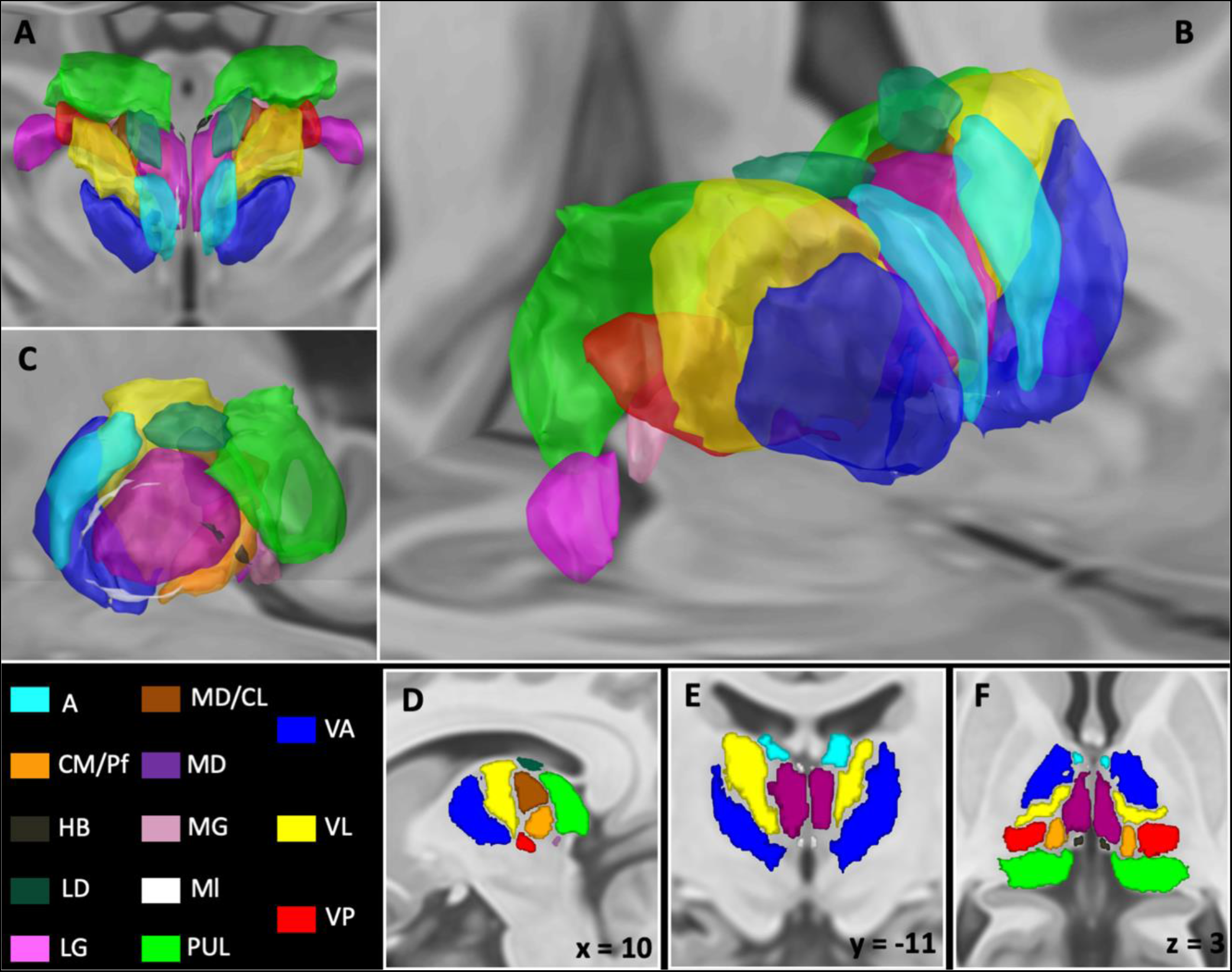
The population template-level thalamic atlas. Thalamic nuclear ROIs manually depicted on the population template have been coregistered to the standard T1 ICBM 2009b brain template. A) 3D rendering, top view; B) Antero-lateral view; C) Medial view; D) 2D single slice, sagittal section; E) coronal section; F) sagittal section. Color labels are described in the figure.

### Similarity analysis

Table 2 shows volumes of thalamic nuclei in mm^3^. In general, the average volume of subject-level thalamic maps is lower than the volume of thalamic maps drawn on the population template, that are, in turn, smaller than those derived from histological maps (Figure 8). Volume estimates of thalamic nuclei are stable across different subjects and population template (ICC=0.996; 95% Confidence Interval=0.993-0.998).

**Table 2.** Volumes of thalamic nuclear maps. Average volumes of thalamic nuclei at subject-level are compared to template-level parcellation and to histological atlas. Volumes are expressed in mm^3^. StD: standard deviation.

|  | Left |  |  | Right |  |  |
| --- | --- | --- | --- | --- | --- | --- |
|  | *Subject-level*<br>*(mean ± StD)* | *Template-*<br>*level* | *Histological* | *Subject-level*<br>*(mean ± StD)* | *Template-*<br>*level* | *Histological* |
| **A** | 332.8 ± 64.7 | 569.1 | 941.1 | 318.4 ± 91.86 | 533.1 | 858.5 |
| **CM/Pf** | 389.4 ± 116.1 | 520.3 | 973 | 392.4 ± 100.3 | 521.5 | 870.6 |
| **Hb** | 33.23 ± 11.64 | 70.25 | 223.1 | 42.54 ± 17.36 | 63.5 | 178.5 |
| **LD** | 242.5 ± 63.16 | 294.5 | 612.9 | 187.4 ± 51.76 | 250.9 | 502.9 |
| **LG** | 321.5 ± 113.5 | 329.4 | 696.6 | 294.5 ± 89.6 | 390 | 595.8 |
| **MD/CL** | 241.5 ± 85.1 | 376.5 | 880.8 | 259.9 ± 50.05 | 267.1 | 779 |
| **MD** | 755.7 ± 140.8 | 1166 | 2309 | 771.4 ± 133 | 1359 | 2164 |
| **MG** | 198.4 ± 75.01 | 242.5 | 608.9 | 179.5 ± 42.57 | 192.5 | 530.3 |
| **MI** | 162.8 ± 82.59 | 108.4 | 1080 | 126.9 ± 71.37 | 157 | 947.8 |
| **Pul** | 2350 ± 154.7 | 2721 | 5357 | 2161 ± 187.3 | 2874 | 4916 |
| **VA** | 1196 ± 226.2 | 2023 | 2341 | 1239 ± 159 | 1921 | 2233 |
| **VL** | 1407 ± 91.4 | 2067 | 3375 | 1326 ± 102.3 | 2010 | 3103 |
| **VP** | 569.2 ± 135.5 | 933.3 | 1576 | 567.4 ± 87.92 | 743.8 | 1393 |

**Figure 8.**
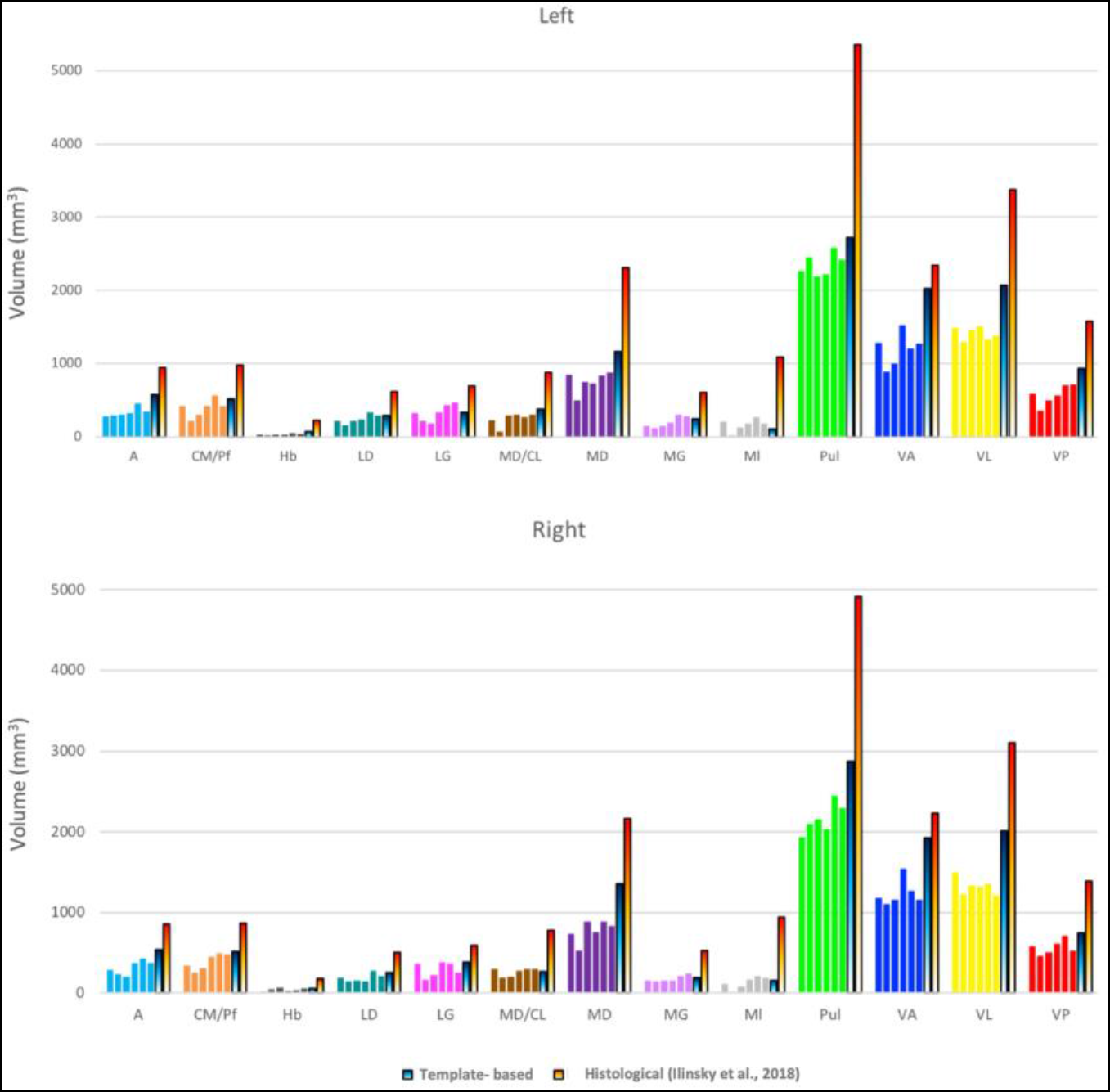
Volumes of thalamic nuclei: comparison to histology. Bar plots illustrating volumes (in mm^3^) of right and left thalamic nuclei in comparison between single-subject estimates (colors), template-level estimates (cold) and histological estimates (hot).

Similarity values of manually-derived thalamic maps are summarized in Table 3. Thalamic maps show OBL values ranging from 0.20 to 0.66 (average: 0.45); the thalamic nuclei scoring the highest OBL values are Pul (left: 0.66; right: 0.64), VA (left: 0.64; right: 0.63) and MD (left: 0.61; right: 0.58), while Ml (left: 0.22; right: 0.2), Hb (left: 0.26; right: 0.21) and LD (left: 0.4; right: 0.28) resulted in the lowest OBL values. The TAO value obtained for the whole parcellation was 0.43.

Template-level thalamic parcellation showed high similarity to histological parcellation, with Dice coefficient values ranging from 0.10 to 0.83 (average: 0.54), with higher Dice for VA (left: 0.8; right: 0.83), A (left: 0.7; Right 0.73) and MD (left: 0.65; right: 0.74) and lower Dice for Ml (left: 0.11; right: 0.19), Hb (left: 0.27; right: 0.29) and LD (left: 0.39; right: 0.46). Subject-level parcellation, by contrast, retrieved lower Dice values (Diceavg ranging from 0.11 to 0.56, average: 0.32), with Pul (left: 0.56; right: 0.55), VA (left: 0.56; right: 0.48) and MD (Right: 0.46; Left: 0.49) scoring the highest values, while Ml (left: 0.12; right: 0.12), LD (left: 0.18; right: 0.18) and LG (left: 0.22; right: 0.14) the lowest values.

**Table 3.** Similarity analysis. Results of similarity measures for inter-subject similarity (OBL) and similarity to histological atlas both for the template-level (Dice) and the subject-level parcellation (Diceavg). L: left. R: right.

|  | OBL |  | Dice<br>(template<br>level) |  | Dice<sub>avg</sub><br>(subject level) |  |
| --- | --- | --- | --- | --- | --- | --- |
|  | L | R | L | R | L | R |
| **A** | 0.5 | 0.45 | 0.7 | 0.73 | 0.3 | 0.33 |
| **CM/Pf** | 0.49 | 0.51 | 0.54 | 0.61 | 0.48 | 0.46 |
| **Hb** | 0.26 | 0.21 | 0.27 | 0.29 | 0.2 | 0.27 |
| **LD** | 0.4 | 0.28 | 0.39 | 0.46 | 0.18 | 0.18 |
| **LG** | 0.39 | 0.5 | 0.56 | 0.63 | 0.22 | 0.14 |
| **MD/CL** | 0.31 | 0.31 | 0.48 | 0.47 | 0.19 | 0.23 |
| **MD** | 0.61 | 0.58 | 0.65 | 0.74 | 0.46 | 0.49 |
| **MG** | 0.39 | 0.37 | 0.5 | 0.47 | 0.17 | 0.2 |
| **MI** | 0.22 | 0.2 | 0.11 | 0.19 | 0.12 | 0.12 |
| **Pul** | 0.66 | 0.64 | 0.65 | 0.65 | 0.56 | 0.56 |
| **VA** | 0.64 | 0.63 | 0.8 | 0.83 | 0.56 | 0.49 |
| **VL** | 0.57 | 0.53 | 0.59 | 0.66 | 0.4 | 0.34 |
| **VP** | 0.49 | 0.46 | 0.63 | 0.54 | 0.42 | 0.37 |

### Connectivity fingerprints of thalamic nuclei

Results of structural and functional connectivity analysis show that, for each of the obtained thalamic nuclei, a well-recognizable, distinct connectivity pattern can be identified. The connectograms in Figure 9 show a quantitative representation of these connectivity fingerprints, while values of structural (F%) and functional (peak t) connectivity for each pair of thalamic nuclei to each target ROI are detailed in Supplementary Results. Average population tractograms and activation clusters for each nucleus are displayed in Supplementary Figures 1-3.

**Figure 9.**
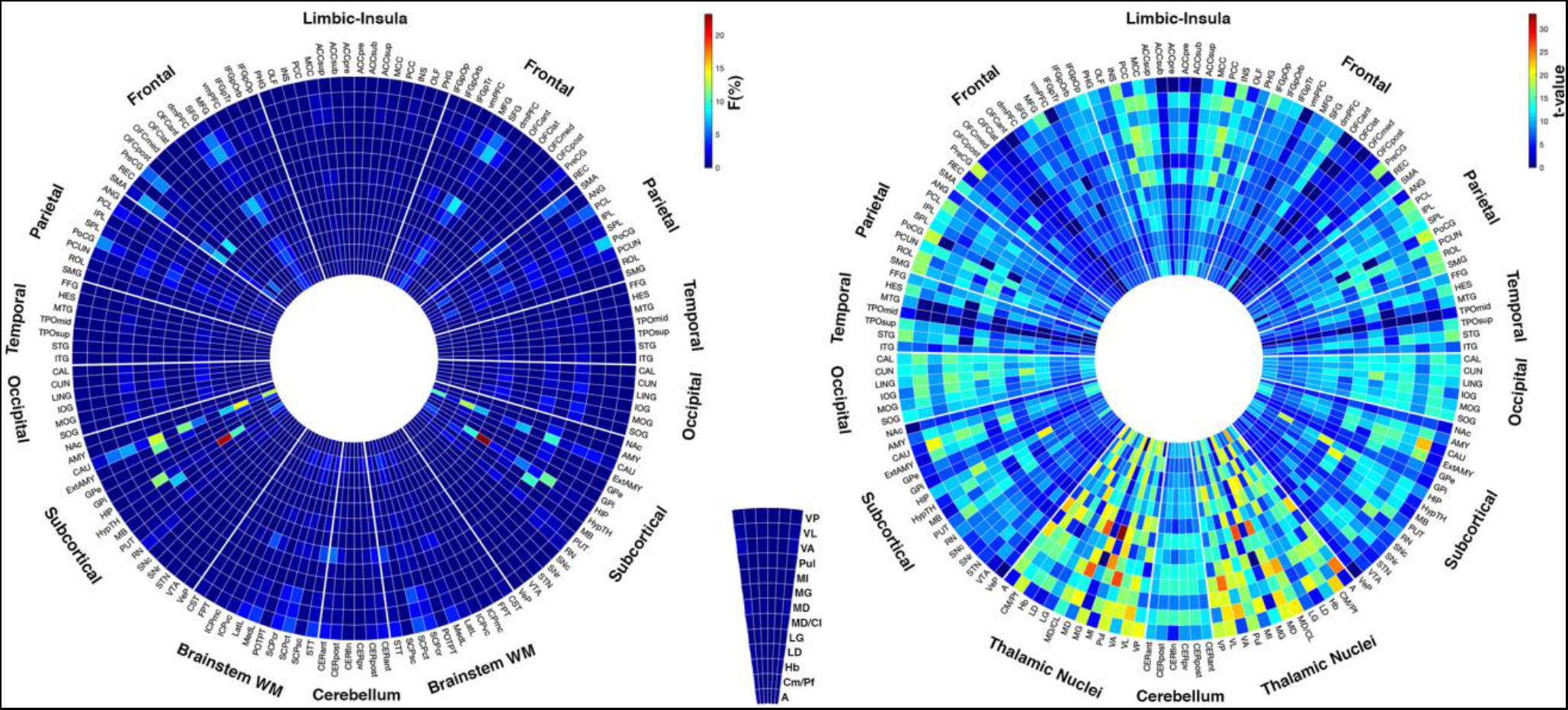
Connectivity fingerprints of thalamic nuclei. Connectograms in form of circular heatmaps illustrating structural (left) and functional (right) connectivity profiles for each thalamic nucleus. Structural connectivity is expressed as percentage of streamline weights (F%), while functional connectivity as peak t-statistic (t-value) values for each target ROI. Left hemispheric targets are depicted on the left, right hemispheric targets on the right, and midline ROIs on the bottom. Abbreviations can be found in the supplementary excel spreadsheet file.

## Discussion

In the present study, we provide high-resolution group-level maps of 13 thalamic nuclei using an optimized short-tracks TDI approach, demonstrating the usefulness of such method for individualized identification of thalamic nuclear areas. To the best of our knowledge, the present study is the first to evaluate the anatomical information of the stTDI approach for human thalamic imaging at 3T field strength.

The high-resolution, high-quality images shown in the current study demonstrate the useful complementary information that can be obtained using super-resolution stTDI. While the feasibility of using stTDI to gain *in vivo* insights into thalamic anatomy has been already suggested from a previous work on 7T MRI datasets (Calamante et al. 2013), the present study pushes forward this concept by describing an optimized pipeline for multi-contrast stTDI mapping and manual delineation of thalamic nuclei at 3T field strength, both on single subjects and on a group template. In line with previous evidence showing a good qualitative agreement between TDI and both myelin- and Nissl-stained histological sections in non-human and human brain (Calamante et al. 2010; Calamante, Tournier, Kurniawan, et al. 2012), the present work shows that matched histological sections can be effectively used to improve the identification of anatomical structures on high-resolution TDI maps.

Our results demonstrate that our protocol for manual delineation of thalamic structures leads to robust thalamic nuclear estimates across different subjects. In particular, volumetric estimates of thalamic nuclei have a very high ICC (0.996, 95% CI=0.993-0.998), suggesting that our proposed manual delineation method could be able to reliably assess thalamic morphometry. In line with this result, manually-delineated thalamic nuclei show also good structural similarity, as suggested by the relatively high TAO value (0.43) for the entire parcellation, and by high OBL values (average: 0.45; range: 0.20-0.66).

When compared to histology, however, it should be noted that both template-level and subject-level parcellations may systematically underestimate thalamic volumes: this result may be explained by taking into account that stTDI cannot identify clear-cut boundaries for thalamic nuclei, leading to very conservative delineations of thalamic substructures. This feature may also explain the differences observed between histological and stTDI-based thalamic maps in terms of structural similarity. Our results show that template-level manual outlines of thalamic nuclei have, in general, a good agreement with histological-derived ROIs, with an average Dice value of 0.54 (range: 0.10-0.83), while average Dice of subject-level thalamic maps show generally lower values. This could be partly explained by the anatomical variability within the human thalamus, considering that the histological atlas is obtained from specimens of a single subject (Ilinsky et al. 2018) and is thus unable to account for inter-subject variability. We suggest that, on thalamic maps obtained from single subjects, the effects of inter-individual differences may strongly affect similarity measures; conversely, group-level thalamic estimates are based on an average of a higher number of subjects (n=100), and may be then more effectively generalized to match the individual anatomy displayed by single-subject histological estimates. Both for template and subject-level parcellation, similarity with their histological-based counterparts was higher for larger nuclei such as VA, A, MD, VL, VP or Pul. While these results may depend on the well-known bias of Dice coefficient when evaluating smaller structures (Shamir et al. 2016), a similar trend was found for inter-subject structural similarity measures (Table 3), that have been corrected for structure volumes: this result may suggest that specific features of stTDI contrast would make these nuclei more easily identifiable, leading to more stable delineations in comparison both to histology and between different subjects. Noteworthy, these nuclei are markedly hypointense, in particular in AFD-stTDI maps, thus showing a good contrast with the surrounding structures and better-defined boundaries. By contrast, other nuclei such as MD/CL, CM/Pf, geniculate nuclei or LD show “fuzzy” and less defined boundaries and, subsequently, lower similarity values; such effects are emphasized for smaller nuclei such as Hb or Ml, which show the lowest similarity values.

Taken together, these results confirm the potential of stTDI approach as a useful additional imaging modality, which can allow for a direct identification of small brain structures, but with the advantages of being applicable *in vivo* to the human brain, and in a potentially individualized way, i.e. tailored to work on a single subject scale. This may help to overcome most of the traditional well-known limitations of histology-based atlases, which include i) the inability to account for anatomical variability (Krauth et al. 2010), ii) the artifacts connected to shrinking and deformation of structures of interest after post-mortem fixation (Chakravarty et al. 2006), and iii) the reliance on an accurate registration to a reference template to properly match anatomical structures of interest when they are non-linearly coregistered to subject images (Ewert et al. 2018). While many other attempts to directly identify thalamic sub-structures *in vivo* using structural MRI have been performed, many of these methods need very long acquisition times (Deoni et al. 2005; Lemaire et al. 2010), are focused on just a few nuclei or nuclear groups (Spiegelmann et al. 2006; Vassal et al. 2012; Erica et al. 2014; Möttönen et al. 2015; Jiltsova et al. 2016) or require ultra-high field 7T MRI scanners with customized, non-standard pulse sequences (Kanowski et al. 2014; Tourdias et al. 2014; Su et al. 2019; Liu et al. 2020). By contrast, the method proposed in the present study is able to provide a fine-grained manual thalamic parcellation, relying mostly on DWI sequences that could be more routinely acquired in a clinical and research context with a 3T MRI scanner.

In the last decades, many individualized multimodal imaging methods for thalamic parcellation have been proposed. In particular, many of these methods are based on the application of clustering algorithms to local features of the structural or DWI MRI signal, both directly or after specific signal modeling (such as FOD modeling), following the assumption that different thalamic nuclei can be distinguished by different signal features (Deoni et al. 2007; Traynor et al. 2011; Battistella et al. 2017; Najdenovska et al. 2018). While many of these methods claim high similarity to histological atlases (i.e Morel atlas), such similarity has been sparsely tested in a quantitative fashion. In addition, the extensive use of clustering algorithms (e.g. k-means), that work better to identify roughly spherical clusters of similar size (Jain 2010; Eickhoff et al. 2015), may lead to an over-simplification of the complex thalamic anatomy.

Other popular methods for parcellation of thalamic structures are based on global, per-voxel estimates of connectivity, measured both with diffusion-based tractography (structural connectivity) (Behrens et al. 2003; Traynor et al. 2010; O’Muircheartaigh et al. 2011; Lambert et al. 2017) or resting-state fMRI (functional connectivity) (Fan et al. 2015; Ji et al. 2016; Kumar et al. 2017; van Oort et al. 2018). The rationale behind these approaches is that voxels belonging to different thalamic nuclei would show distinctive connectivity features. Being this assumption not entirely true, as different nuclei could share similar connectivity profiles, parcellations derived from these methods could be more properly interpreted as identifying coherent, topographically organized functional territories within the human thalamus, rather than nuclei itself (Eickhoff et al. 2015); nevertheless, some of these parcellations have shown some degrees of similarity to histological parcellation (Lambert et al. 2017) and may have clinical utility in contexts in which the individuation of a specific connectivity-defined territory, rather than a proper histological nucleus, is required (Akram et al. 2018; Middlebrooks et al. 2018; Cacciola, Milardi, et al. 2019; Bertino et al. 2020).

Despite that, these methods are conditioned by the well-known limitations concerning the interpretation of streamline density (for tractography-based methods) and between-voxel correlation/covariance of blood oxygen level dependent (BOLD) signal (for resting-state fMRI) as measures of neural connectivity (Buckner et al. 2013; Jones et al. 2013). In addition, they can be strongly influenced by different pre-processing and signal modeling pipelines, with lack of a recognized “gold standard” procedure (Maier-Hein et al. 2017; Parkes et al. 2018).

The method proposed in the present work is based on tractography and is then conditioned by most of the common drawbacks of this technique (motion, geometrical and susceptibility-induced artifacts, dependence of results from the signal modeling approach employed) (Calamante 2017). However, in this case, tractography is not employed to quantify neural “connectivity”, but rather to provide improved anatomical contrast and super-resolution features. Furthermore, the use of short-tracking makes tractographic reconstruction more sensitive to local (instead of global) features of the diffusion signal and ensures a total independency of stTDI contrast from length and distance biases, very common limitations of diffusion tractography (Dhollander et al. 2014).

Above and beyond reconstructing maps of the human thalamic nuclei by employing an optimized super-resolution stTDI approach, the parcellation established in the present study can be of great utility both for neuroimaging studies as well as stereotactic targeting for functional neurosurgery. Hence, we have made publicly available all the thalamic maps obtained by manual segmentation of the population template-based stTDI (https://github.com/BrainMappingLab), non-linearly coregistered to the most updated MNI template (ICBM 2009c nlin asym) (Fonov et al. 2009).

### Structural and functional connectivity fingerprints of human thalamic nuclei

In order to demonstrate the potential usefulness of the proposed approach in a neuroimaging research context, we coregistered the group-level thalamic maps on 210 healthy subjects from the HCP repository and performed seed-based structural and functional connectivity analysis thus obtaining specific thalamic connectivity profiles. To our knowledge, only a few works describe thalamic connectivity at the nuclear level with such detail. Lambert and colleagues (Lambert et al. 2017) after thalamic parcellation based on probabilistic tractography distribution feature maps, described whole-brain structural connectivity profiles for each of the obtained thalamic nuclei. Our results widen the perspective on thalamic nuclei connectivity by including a more detailed thalamic parcellation (13 nuclei vs 9), a more granular cortical (Rolls et al. 2020) and subcortical (Pauli et al. 2018) parcellation and additional information on connectivity with brainstem white matter tracts (Tang et al. 2018). In line with their findings, and with experimental results in different animal species, we report high structural connectivity to caudate nucleus, extended amygdala, hippocampus, hypothalamus and frontal lobe regions for the A nuclei (Child and Benarroch 2013), while LD nuclei showed high connectivity to precuneus, superior parietal and frontal cortex, occipital lobes, striatum and hippocampus (Mizumori and Williams 1993; van Groen et al. 2002; Bezdudnaya and Keller 2008). Among intralaminar nuclei, we found high structural connectivity to prefrontal cortex, caudate and hippocampus for Ml nuclei, primary sensorimotor cortex, prefrontal cortex and ascending brainstem tracts for CM/Pf, and sensorimotor parietal cortex, in particular for MD/CL, in line with previous findings (Saalmann 2014). We report also high structural connectivity to striatum and prefrontal and sensorimotor cortices for thalamic nuclei of the ventral group; in particular, VA shows higher structural connectivity to prefrontal cortices, and VL to sensorimotor cortical areas, in line with their differential roles in the cortico-thalamic-basal ganglia circuitry (McFarland and Haber 2002; DeLong and Wichmann 2010); on the other hand, VP shows high structural connectivity to sensorimotor frontal and parietal cortices such as precentral and postcentral gyrus, and to brainstem ascending pathways involved in somatic sensation, including medial lemniscus and spinothalamic tract (Padberg et al. 2009). The MD nuclei also exhibited high structural connectivity to striatum, prefrontal, premotor and cingulate cortices, as suggested by most of the existing literature (Mitchell and Chakraborty 2013; Parnaudeau et al. 2018). We also describe dense structural connectivity for Pul to occipital visual regions and to parietal and prefrontal cortices, in keeping with previous investigations (Leh et al. 2008; Benarroch 2015), while the metathalamic nuclei (LG and MG) show similar connectivity profiles involving mostly occipital and temporal regions and hippocampus. Noteworthy, we were not able to find high structural connectivity between MG and primary auditory cortex (acoustic radiations), as it would be expected: this result could be conditioned by the peculiar course of these connections, that makes them very difficult to be reconstructed using tractography (Maffei et al. 2019); despite that, we report higher connectivity for MG in comparison to LG to the lateral lemniscus, that is part of the ascending auditory pathway (Sitek et al. 2019). Finally we described structural connectivity profiles of Hb, that shows high connectivity to prefrontal cortex, basal ganglia, superior cerebellar peduncle and subcortical structures including amygdala and hypothalamus, in keeping with both human and non-human investigations (Herkenham and Nauta 1979; Shelton et al. 2012; Strotmann et al. 2014; Quina et al. 2015).

Beyond a quantitative and qualitative description of structural connectivity profiles of thalamic nuclei, the present work adds useful insights on thalamic functional connectivity. Our results, similarly to previous investigations (Zhang et al. 2010), show substantial degree of concordance between functional and structural connectivity profiles of thalamic nuclei; on the other hand, while structural connectivity profiles are inherently constrained by the reconstruction of direct anatomical connections only, functional connectivity profiles revealed, for each thalamic nucleus, distinct but partially overlapping, widespread connectivity to cortical, subcortical and cerebellar regions. These results are in keeping with other recent findings suggesting that thalamic nuclei show functional connectivity to areas involved in multiple cortical networks, and may play an integrative role rather than simply relaying information from and to the cerebral cortex (Kultas-Ilinsky et al. 2003; Hwang et al. 2017). In line with this view, we also report strong functional connectivity between each thalamic nucleus and other thalamic nuclei, suggesting the presence of strong co-activation patterns between distinct thalamic nuclei. These coactivation patterns may contribute to the integration of cortical information across distinct brain networks and reflect a hierarchical flowing of information across corticothalamic functional connectivity, similar to what has been described in a recent work employing connectivity gradient embedding (Yang et al. 2020).

### Implications for stereotactic targeting in functional neurosurgery

These results suggest that the thalamic maps obtained with template level manual parcellation could be useful in the neuroimaging field to further improve our knowledge on the thalamic connectome in healthy individuals. In addition, these maps may also help to better elucidate the role of thalamic substructures involved in different brain diseases, such as MS (d’Ambrosio et al. 2017; Lin et al. 2019), psychosis (Paez et al. 2011; Anticevic et al. 2014; Murray and Anticevic 2017; Ramsay 2019), PD, (Planetta et al. 2013; Owens-Walton et al. 2019), AD and cognitive impairment (Wang et al. 2012; Zhou et al. 2013; Cai et al. 2015), migraine (Granziera et al. 2014; Amin et al. 2018) and many other neuropsychiatric conditions (Yin et al. 2011; Banks et al. 2016; Preller et al. 2018). Finally, the present thalamic parcellation could also be useful in the functional neurosurgery setting, as it includes many of the main thalamic targets for DBS and stereotactic ablation. Specifically, the inferior portion of VL nucleus (ventral intermediate, VIM, in Hassler’s nomenclature) (Hassler 1959), is commonly targeted in PD and essential tremor (Cury et al. 2017; Fiechter et al. 2017); the anterior nuclear group is increasingly used in treatment-resistant epilepsy (Fisher et al. 2010), along with CM/Pf (Son et al. 2016), that is also ablated to treat chronic central pain (Young et al. 1994); finally, both CM/Pf and MD nuclei have been proposed as targets for treatment of GTS (Vandewalle et al. 1999; Vernaleken et al. 2009; Ackermans et al. 2011).

The thalamic delineation protocol proposed in the present paper is able to directly identify VL, CM/Pf, MD and anterior nuclei with good-to-high reliability (OBL values ~ 0.5) and good similarity to histological ground-truth.

On the other hand, a crucial topic in neurosurgical literature is *in vivo* identification of the VIM nucleus, which is likely to be ideally located at the entrance area of pre-lemniscal radiation coming from the cerebellum (Hassler 1959, 1983). While currently the debate on the precise histological identity of this structure is still open (Mai and Majtanik 2019), histological, clinical and neuroimaging evidence suggest that it may be located within the inferior portion of VL nucleus (Cury et al. 2017; Akram et al. 2018; Ilinsky et al. 2018). Hence, the putative position of VIM may be inferred from our parcellation by projecting the anterior commissure-posterior commissure (AC-PC) plane on the VL map, as suggested by different authors (Ilinsky et al. 2018; Su et al. 2019)

Non-linear registration of thalamic maps to subject space may represent a valuable option in clinical settings when good-quality DWI are not available. However, when feasible, the protocol for individualized TDI reconstruction proposed in the present work could be employed to get a more precise and patient-tailored target identification. In addition, such approach could provide further information also on some extra-thalamic structures of known surgical interest, such as the ansa lenticularis, the zona incerta or the mammillothalamic tract (Rossi et al. 2016; Balak et al. 2018; Ossowska 2020).

### Methodological considerations and conclusions

While our results have been obtained on high quality 3T DWI datasets from the HCP repository employing a multi-shell multi-tissue CSD approach for FOD estimation, which relies on accurate T1-DWI coregistration and DWI geometrical distortion correction (Jeurissen et al. 2014b), it can be hypothesized that the use of single-shell CSD to estimate multi-tissue FOD maps could make the process entirely depending on the DWI dataset alone (Dhollander et al. 2016).

In addition, the fact that the analysis is limited to the diencephalic region, that is usually less affected from DWI geometrical distortions (Treiber et al. 2016), could reduce the dependency of results on diffusion preprocessing techniques and speed up the acquisition process by reducing the field of view. However, we acknowledge that further investigations are needed to evaluate the effects of preprocessing, signal modeling and tractography parameters on stTDI contrast and thalamic delineation. Furthermore, while manual delineation is time-consuming and requires further knowledge on thalamic anatomy, the parcellation process could be fully automatized in the future, by using a multi atlas approach based on label fusion (Chakravarty et al. 2013; Hongzhi Wang et al. 2013; Iglesias et al. 2018b; Su et al. 2019; Liu et al. 2020)

In the present work, we devised a protocol for the identification of thalamic nuclei based on stTDI imaging. We showed that such approach can accurately identify thalamic sub-structures in good agreement with histological ground-truth, and with high inter-subject reliability. In addition, we provided a group-level atlas of thalamic nuclei in a commonly used template space. We also employed this atlas in order to comprehensively reconstruct thalamic structural and functional connectivity profiles in a large cohort of healthy subjects.

We hope that the proposed parcellation protocol and group-level thalamic atlas can be of interest both for basic and translational neuroimaging studies and clinical applications in functional neurosurgery.

## Acknowledgments

The histological images reproduced in this paper were obtained from http://www.humanmotorthalamus.com/ courtesy of prof. Igor Ilinsky and prof. Kristy Kultas-Ilinsky. To them go also our most sincere thanks for their kind and valuable help during the preparation of the present work, and their useful suggestions and constructive critique on the manuscript.

Data used in this paper were provided by the Human Connectome Project, WU-Minn Consortium (Principal Investigators: David Van Essen and Kamil Ugurbil; 1U54MH091657). The Human Connectome Project was funded by the 16 NIH institutes and centers that support the NIH Blueprint for Neuroscience Research; and by the McDonnell Center for Systems Neuroscience at Washington University. This research did not receive any specific grant from funding agencies in the public, commercial, or not-for-profit sectors.

## Supplementary Results

### Structural and functional connectivity profiles of thalamic nuclei

Structural and functional connectivity profiles of each thalamic nucleus to all the cortical and subcortical targets employed are included in a supplementary excel spreadsheet file. In the present document we focus only on the highest values of structural (F) and functional (peak-t) connectivity for each nucleus.

#### Anterior nuclei (A)

The A nuclei showed high structural connectivity to left (F=1.07%) and right (F=1.79%) pars triangularis (IFGpTr), left (F=2.94%) and right (F=2.4%) middle frontal gyri (MFG), left (F=4.92%) and right (F=3.89%) superior frontal gyri (SFG), left (F=2.98%) and right (F=3.92%) dorsomedial prefrontal cortex (dmPFC), left (F=0.92%) and right (F=1.31%) supplementary motor areas (SMA) and left (F=0.96%) and right (F=1%) precuneus (PCUN); among subcortical structures, high structural connectivity was found for bilateral caudate (CAU) (left: F=10.26%; right: F=12.61%), hippocampus (HIP) (left: 4.44%; right: 5.35%), hypothalamus (HypTH) (left: F=3.88%; right: F=2.83%) and left extended amygdala (ExtAMY) (F=4.1%). Concerning functional connectivity, high peak-t values were found in bilateral cingulate cortex: pregenual anterior cingulate (ACCpre), (left: t=10.2; right: t=9.87); supracallosal anterior cingulate (ACCsup) (left: t=11.5; right: t=10.41); middle cingulate cortex (MCC) (left: t=12.39; right:12.73) and posterior cingulate cortex (PCC) (left: t=14.1; Right: t=12.93). High functional connectivity was also found to left pars opercularis (IFGpOp) (t=12.39), left (t=10.87) and right (11.45) PCUN, and bilateral occipital lobe: calcarine cortex (CAL) (left: t=9.85; Right: t=10.66), cuneus (CUN) (left: t=9.54; right: t=11.08), lingual gyrus (LING) (left: t=9.25; Right: t=12.36), inferior (IOG) (left: t=9.85; right: t=10.66) and middle occipital gyri (MOG) (left: t=10.33; Right: t=10.43). Among subcortical structures, high values were also obtained for amygdala (AMY) (left: t=11.05; right: t=11.9), CAU (left: t=13.53; right: t=14.8), ExtAMY (left: t=11.76; right: t=13.3), putamen (PUT) (left: t=10.8; right: t=14.38) while among thalamic nuclei, Hb (left: t=23.31; right: t=18.79), right LD (t=15.76), MD/CL (left: t= 10.29; right: t=10.45), MD (left: t=21.43; right: t=20.66), Ml (left: t=12.95; right: t=19.13), Pul (left: t=16.27; right: t=15.69), right VA (t=19.07), and VL (left: t=14.23; right: t=18.03) scored high peak t-values. High functional connectivity was also found bilaterally in the posterior hemispheres of the cerebellum (CERpost) (left: t=10.37; right: t=10.37) and in the posterior vermis (CERpv) (t=10.65).

#### Centromedian/parafascicular complex (CM/Pf)

High structural connectivity was found for CM/Pf, to bilateral MFG (left: F=1.32%; right: F=1.37%), SFG (left: F=2.95%; right: F=3.83%), precentral gyrus (PreCG) (left: F=3.48% ; right: F=5.52%), right dmPFC (F=1.01%), SMA (left: F=2.96% ; right: F=4.54%), paracentral lobule (left: F=4.52%; right: F=1.58%), postcentral gyrus (PoCG) (left: F=4.31%; right: F= 2.31%) and left PCUN (F=1.04%). For subcortical structures, relatively high structural connectivity values were obtained to left (F=2.10%) and right (F=3.80%) CAU, medial lemniscus (MedL) (left: 2.25%; right: 3.08%), right lateral lemniscus (LatL) (F=2.01%), left (F=1.97%) and right (F=3.7%) spinothalamic tracts, and both cerebello-thalamic and cerebello-rubral divisions of the superior cerebellar peduncle (SCPct, left: F=3.86%, right: F=3.7%; SCPcr, left: F=3.08%, right=3.58%), together with anterior cerebellum (CERant) (left: F=2.19%; right: F=2.24%) and CERpost (left: 1.38%; right: 1.46%).

Target cortical regions obtaining high peak-t values were left (t=9.63) and right (t=9.71) MCC, left (t=10.84) and right (t=9.41) insula (INS), left (t=9.64) dmPFC, right PCUN (t=9.6) and left (t=10.85) and right (t=9.41) superior temporal gyri (STG). High functional connectivity was also obtained to thalamic nuclei such as left A (t=18.87), Hb (left: t=9.51; right: t=14.67), left LD (t=20.06), MD/CL (left: t=19.23; right: t=17.49), MD (left: t=19.16; right: t=18.71), Pul (left: t=15.36; right: t=16.37), VL (left: t=18.09; right: t=16.14) and right VP (t=16.2).

#### Habenula (Hb)

The Hb shows high structural connectivity to left IFGpOrb (F=1.57%), left (F=1.71%) and right (F=1.03%) IFGpTr, left (F=3.28%) and right (F=2.53%) MFG, left (F=5.66%) and right (F=4.66%) SFG, left (F=3.05%) and right (F=3.87%) dmPFC and left (F=1.49%) and right (F=2.34%) SMA. Subcortical structures showing high structural connectivity to Hb were left AMY (F=1.07%), and bilateral CAU (left: F=12.44%; right: F=14.17%), extAMY (left: F=1.32%; right: F=3.86%), left external globus pallidus (GPe) (left: F=1.28%;), HypTH (left: F=3.61%; right: F=1.4%) and PUT (left: F=3.13%; right: F=1.33%). Brainstem white matter tracts such as left MedL (F=1.17%), left (F=1.05%) and right (F=1.27%) SCPcr, left (F=2.09%) and right (F=1.78%) SCPct, and right STT (F=1.09%) obtained relatively high structural connectivity values.

Higher peak-t values were found bilaterally for MCC (left: t=8.23; right: t=8.74), PCC (left: t=8.9; right: t=8.11), CAL (left: t=8.80; right: t=10.66), LING (left: t=8.80; right: t=8.21), CAU (left: t=7.8; right: t=8.14) and HIP (left: t=18.9; right: t=8.28). Among thalamic nuclei, high functional connectivity was found to right CM/Pf (t=11.94), left (t=19.4%) and right (t=8.27) LG, right (t=8.44) MD/CL, left (t=13.78) and right (t=19.93) MD, left MG (t=16.32), left (t=16.11) and right (t=19.21) Pul, left (t=9.15) and right (t=11.52) VL, and left VP (t=8.81). For cerebellar cortex, the flocculonodular lobe (CERfln) (t=9.58), CERpv (t=8.98), and bilateral CERpost (left: t=8.98; right: t=8.59).

#### Lateral dorsal nuclei (LD)

The LD nuclei showed high structural connectivity to left IFGpTr (F=1.05%), MFG (left: F=2.09%; right: F=2.16%), SFG (left: F=3.49%; right: F=3.85%), dmPFC (left: F=1.74%; right: F=2.23%), SMA (left: F=1.62%; right: F=3.34%), left inferior parietal lobule (IPL) (F=1.1%), superior parietal lobule (SPL) (left: F=2.04%; right: F=1.63%), PoCG (left: F=1.31%; right: F=1.74%), PCUN (left: F=2.12%; right: F=1.94%) and left MOG (F=1.28%). Subcortical structures showing high connectivity to LD were left (F=7%) and right (F=7.55%) CAU, left (F=9.61%) and right (F=10.13%) HIP, left HypTH (F=1.57%), and left PUT (F=1.10%).

Cortical areas showing high functional connectivity to LD include MCC (left: t=10.38; right: t=11.9), PCC (left: t=12.33; right: t=12.33), left dmPFC (t=9.4) SMA (left: t=9.43; right: t=9.43), PCUN (left: t=10.98; right: t=10.86), CAL (left: t=10.46; right: t=11.71), CUN (left: t=9.37; right: t=10.86), LING (left: t=9.59; right: t=11.71); IOG (left: t=10.45; right: t=11.71), MOG (left: t=9.83; right: t=10.5) and left SOG (t=10.03). High peak-t values were found also for subcortical regions such as left (t=11.28) and right (t=8.7) AMY, left (t=10.65) and right (t=11.6) CAU, left (t=11.28) and right (t=10.24) PUT, and, among thalamic nuclei, A (left: t=9.82; right: t=19.76), left CM/Pf (t=17.47), Hb (left: t=11.23; right: t=10.69), MD/CL (left: t=21.23; right: t=13.37), MD (left: 20.54; right: t=16.49), right ML (t=10.09), Pul (left: t= 18.34; right: t=18.05), right VA (t=10.01), VL (left: t=21.23; right: t=16.03) and left VP (left: t=16.44). High functional connectivity was also found to cerebellar cortex, in particular CERpost (left: t=10.8; right; t=10.8) and CERpv (t=10.8).

#### Lateral geniculate nuclei (LG)

The LG nuclei showed high structural connectivity to left SPL (F=1.17%), PCUN (left: F=1.5%; right: F=1.8%) and to temporal and occipital cortical regions, such as middle temporal gyrus (MTG) (left: F=1.14%; right: F=1.31%), middle temporal portion of the temporal pole (TPOmid) (left: F=1.18%; right: F=1.24%), inferior temporal gyrus (ITG) (left: F=1.74%; right: F=1.27%), CAL (left: F=1.41%; right: F=1.88%), CUN (left: F=0.98%; right: F=1.48%), LING (left: F=2.95%; right: F=2.95%), MOG (left: F=2.76%; right: F=1.80%) and SOG (left: F=1.72%; right: F=1.87%). LG showed also high structural connectivity to right AMY (F=1.24%), and left (F=23.2%) and right (F=22.04%) HIP.

High peak-t values were found in different cortical regions, including ACCsup (left: t=10.03; right: t=10), MCC (left: t=11.10; right: t=10.78), PCC (left: t=11.6; right: t=10.95), INS (left: t=12.55; right: t=11.95), right parahippocampal gyrus (PHG) (t=10.84), left dmPFC (t=10.59), PCUN (left: t=10.04; right: t=10.33), rolandic operculum (ROL) (left: t=11.65; right: t=10.13), Heschl’s gyrus (HES) (left: t=11.33; right: t=10.84), STG (left: t=12.55; right: t=11.22), CAL (left: t=12.82; right: t=14.25), CUN (left: t=9.9; right: t=10.1), LING (left: t=; right: t=17.8), IOG (left: t=11.4; right: t=11.04) and left MOG (t=10.85). High functional connectivity to LG was also found for subcortical structures, such as left (t=10.89) and right (t=10.7) CAU, left (t=11.17) and right (t=21.11) HIP, left HypTH (t=10.32), right PUT (t=10.93) and left substantia nigra pars reticulata (SNr) (t=10.32). Among thalamic nuclei, LG showed also high functional connectivity to A (left: t=9.8; right: t=14.25), right CM/Pf (t=14.3), Hb (left: t= 10.61; right: t=17.53), MD/CL (left: t=12.2; right: t=17.42), MD (left: t=13.65; right: t=19.88), Pul (left: 14.42; right: t=14.27), VA (left: t=10.03; right: t=13.02) and VL (left: t=13.25; right: t=16.9). LG obtained also high peak-t values to cerebellar regions, including CERpost (left: t=10.96; right: t=10.96) and CERpv (t=10.8).

#### Mediodorsal/centrolateral complex (MD/CL)

High structural connectivity values to MD/CL were obtained from left (F=2.07%) and right (F=1.08%) MCC, left (F=4.31%) and right (F=1.76) MFG, left (F=4.31%) and right (F=5.58%) SFG, left (F=3.87%) and right (F=8.3%) PreCG, left (F=4.74%) and right (F=7.5%) SMA, left (F=4.73%) and right (F=2.08) PCL, left SPL (F=1.3%), left (F=4.14%) and right (F=3%) PoCG, and left PCUN (F=1.52%). MD/CL showed also high connectivity to CAU (left: F=3.24%; right: F=5.28%), left HIP (F=2.54%), left PUT (F=1%), and to brainstem white matter tracts such as right MedL (F=1.37%), right STT (F=1.26%), right (F=1.09%) and left (F=0.97%) SCPcr, right (F=1.53%) and left (F=1.38%) SCPct.

Cortical regions with high functional connectivity to MD/CL include INS (left: t=11.75; right: t=13.22), PreCG (left: t=9.81; right: t=10.2), PoCG (left: t=10.2; right: t=9.82); ROL (left: t=11.75; right: t=12.54), supramarginal gyrus (SMG) (left: t=10.15; t=11.82), HES (left: t=10.86 ; right: t=10.31), STG (left: t=11.56; right: t=10.82), CAL (left: t=9.99; right: t=10.71), and LING (left: t=11.21; right: t=10.78). High peak-t values were obtained also for bilateral HIP (left: t=19.51; right: t=10.17), and left STN (t=9.96). Thalamic nuclei with high functional connectivity values include CM/Pf (left: t=14.16; right: t= 19.69), right LD (t=13.98), left LG (t=18.7), MD (left: t=10.96; right: t=19.12), left MG (t=19.93), Pul (left: t=19.65; right: t=18.57), VL (left: t=10.55; right: t=19.44) and VP (left: t=12.48; right: t=17.99)

#### Mediodorsal nuclei (MD)

The MD shows high structural connectivity to limbic and frontal lobe regions such as right ACCsup (F=1.22%), right MCC (F=1.20%), left IFGpOrb (F=1.17%), IFGpTr (left: F=2.5%; right: F=1.45%), MFG (left: F=4.76%; right: F=4.11%), SFG (left: F=8.35%; right: F=7.23%), dmPFC (left: F=4.09%; right: F=5.19%), right PreCG (F=1.09%), and SMA (left: F=4.12%; right: F=5.3%). MD obtained also high structural connectivity values to CAU (left: F=7.10%; right: F=11.28%), right extAMY (F=2.05%), HIP (left: F=1.82%, right: F=1.01%), left HypTH (F=1.38%) and left PUT (F=2.18%).

High functional connectivity to MD was found for limbic regions such as ACCpre (left: t=12.58; right: t=11.59), ACCsup (left: t=12.58; right: t=12.57), MCC (left: t=12.88; right: t=15.90), PCC (left: t=17.2; right: t=16.78), to left dMPFC (t=12.55), left SPL (t=11.6), left PCUN (t=14.59) and to occipital regions, including CAL (left: t=11.76; right: t=13.36), CUN (left: t=13.05; right: t=14.3), LING (left: t=11.06; right: t=14.21), IOG (left:t=12.15; right: t=13.36), MOG (left: t=12.02; right: t=11.8) and left SOG (t=12.52). The MD shows also high functional connectivity to subcortical structures, including AMY (left: t=13.14; right: t=11.08), CAU (left: t=14.47; right: t=16.45), extAMY (left: t=13.85; right t=12.72), HIP (left: t=10.41; right: t=11.38), HypTH (left: t=20.15; right: t=12.06), mammillary bodies (MB) (left: t=11.81; right: t=12.55), PUT (left: t=13.14; right: t=13.29) and SNr (left: t=11.81; right: t=9.54). Among thalamic nuclei, high peak-t values were found for A (left: t=22.1; right: t=22.66), CM/Pf (left: t=15.52; right: t=17.73), Hb (left: t=15.16; right: t=20.98), LD (left: t=10.22; right: t=15.88), MD/Cl (left: t=11.74; right: t=19.18), Ml (left: t=26.59; right: t=33.25), Pul (left: t=13.49; right: t=20.48), VA (left: t=10.25; right: t=15.6), and VL (left: t=15.74; right: t=17.22). In addition, MD nuclei show also high connectivity to the cerebellar cortex: CERant (left: t=12.55; right: t= 9.96), CERpost (left: t=13.55; right: t=13.55), CERfln (t=12.81) and CERpv (t=13.55).

#### Medial geniculate nuclei (MG)

Cortical regions that obtained high structural connectivity values to MG nuclei include right PreCG (F=1.35%), left IPL (F=1.33%), left (F=2.79%) and right (F=2.48%) SPL, left (F=2.21%) and right (F=4.09%) PoCG, left (F=2.9%) and right (F=2.83%) PCUN, left MTG (F=1.04%), left CAL (F=1.25%), left CUN (F=1.2%), left (F=1.39%) and right (F=2.21%) LING, left MOG (F=3.27%) and left (F=2.11%) and right (F=1.04%) SOG. MG nuclei show also high structural connectivity to HIP (left: F=5.59%; right: F=1.05%), to LatL (left: F=2.34%; right: F=4.72%), MedL (left: F=1.85%; right: F=1.64%), SCPcr (left: F=1.22%; right: F=1.29%), SCPct (left: F=2.63%; right: F=1.52%), STT (left: F=4.23%; right: F=4.47%) and CERant (left: F=2.62%; right: F=5.40%).

The MG nuclei show high functional connectivity to widespread cortical areas, including MCC (left: t=12.1; right: t=11.2), right PCC (t=10.06), INS (left: t=15.12; right: t=17.12%), PHG (left: t=13.9; right: t=10.79), left IFGPOp (t=10.42), %), right MFG (t=10.21), PreCG (left: t=13.32; right: t=13.4), SMA (left: t=12.1; right: t=10.96), PCL (left: t=12.04; right: t=12.34), SPL (left: t=10.92; right: t=10.2), PoCG (left: t=14.22; right: t=12.52), PCUN (left: t=12.91; right: t=11.02), ROL (left: t=15.12; right: t=15.57), SMG (left: t=12.99; right: t=14.52), fusiform gyrus (FFG) (left: t=10.40; right: t=11.65), HES (left: t=13.74; right: t=14.14), superior temporal portion of the temporal pole (TPOsup) (left: t=10.65; right: t=10.7), STG (left: t=12.99; right: t=14.23), CAL (left: t=13.07; right: t=13.11), CUN (left: t=12.41; right: t=10.94), LING (left: t=17.18; right: t=16.1), MOG (left: t=10.13; right: t=9.99) and SOG (left: t=12.41; right: t=10.93). High peak-t values were found also for left CAU (t=10.8), left (t=15.11) and right (t=13.96) HIP, and for the following thalamic nuclei: CM/Pf (left: t=11.61; right: t=19.08), left Hb (t=11.66), MD/Cl (left: t=14.78; right: t=9.85), MD (left: t=14.14; right: t=10.5), Pul (left: t=17.77; right: t=21.75), VL (left: t=13.75; right: t=9.65) and VP (left: t= 14.11; right: t=18.73). In addition, MG nuclei showed high functional connectivity to CERant (left: t=11.65; right: t=11.65) and CERpost (left: t=10.02; right: t=9.66).

#### Midline nuclei (MI)

The MI nuclei show prominent structural connectivity to frontal lobe regions such as MFG (left: F=0.91%; right: F=1.02%), SFG (left: F=1.56%; right: F=1.7%) and right dmPFC (F=1.29%), and to subcortical structures, such as left AMY (F=3.26%), CAU (left: F=9.74%; right: F=13.31%), extAMY (left: F=2.29%; right: F=11.1%), right GPe (F=2.24%), HIP (left: F=2.77%; right: F=1.39%), HypTH (left: F=9.9%; right: F=7.43%) and PUT (left: F=2.23%; right: F=1.94%).

High peak-t values were obtained for ACCpre (left: t=11.55; right: t=10.75), ACCsup (left: t= 11.79; right: t=11.07), MCC (left: t=13.26; right: t=15.82), PCC (left: t=16.65; right: t=16.65), left dmPFC (t=11.86), left SPL (t=12.08), PCUN (left: t=14.61; right: t=13.57), FFG (left: t=10.57; right: t=10.27), and occipital lobe regions such as CAL (left: t=11.01; right: t=13.41), CUN (left: t=12.58; right: t=13.41), LING (left: t=10.76; right: t=13.57), IOG (left: t=12.71; right: t=13.41), MOG (left: t=12.03; right: t=13.25) and SOG (left: t=13.58; right: t=11.05). Ml nuclei also showed high functional connectivity to distinct subcortical structures, including AMY (left: t=11.03; right: t=9.85), CAU (left: t=15.5; right: t=15.2), extAMY (left: t=15.01; right: t=11.7), left GPe (t=13.78HIP (left: t=11.91; right: t=10.32), HypTH (left: t=11.91; right: t=10.27), MB (left: t=10.54; right: t=11.22), PUT (left: t=10.96; right: t=12.49), left SNr (t=10.54) and left STN (t=13.05). Among thalamic nuclei, high peak-t values were obtained for A (left: 17.5; right: 21.73), right CM/Pf (t=15.8), Hb (left: t=12.87; right: t=17.38), left MD/CL (t=10.66), MD (left: t=18.24; right: t=20.64), Pul (left: t=12.24; right: t=16.2), VA (left: t=19.65; right: t=12.16) and VL (left: t=17.41; right: t=12.28). High connectivity was found also to cerebellar cortical regions: CERant (left: t=12.8; right: t=10.49), CERpost (left: t=13.61; right: t=13.61), CERfln (t=13) and CERpv (t=13.61).

#### Pulvinar (Pul)

High structural connectivity values to Pul were found for left (F=1.08%) and right (F=1.39%) SFG, right PreCG (F=1.23%), right SMA (F=1.55%), right angular gyrus (ANG) (F=1.1%), left (F=2.65%) and right (F=2.36%) SPL, left IPL (F=1.57%), left (F=1.62%) and right (F=2.26) PoCG, left (F=3.14%) and right (F=3.37%) PCUN, left (F=1.32%) and right (F=1.19%) MTG, left ITG (F=1.47%), left (F=1.72%) and right (F=1.65%) CAL, left (F=1.27%) and right (F=1.4%) CUN, left (F=3.24%) and right (F=2.23%) LING, left (F=3.29%) and right (F=1.6%) MOG and left (F=2.16%) and right (F=1.73%) SOG. In addition, Pul shows also high structural connectivity to CAU (left: F=1.22%; right: F=2.27%) and HIP (left: F=11.03%; right: F=11.85%).

For what concerns functional connectivity, Pul shows high connectivity to widespread cortical regions, including ACCpre (left: t=12.07; right: t=10.26), ACCsup (left: t=12.84; right: t=12.62), MCC (left: t=13.66; right: t=15.31), PCC (left: t=18.2; right: t=18.19), INS (left: t=13.7; right: t=14.07), left olfactory cortex (OLF) (t=11.07), PHG (left: t=11.14; right: t=11.02), dmPFC (left: t=13.66; right: t=10.89), SMA (left: t=12.94; right: t=10.48), PCL (left: t=11.4; right: t=11.91), left SPL (t=12.9), right IPL (t=10.7), PCUN (left: t=14.86; right: t=14.62), ROL (left: t=11.36; right: t=10.85), SMG (left: t=10.04; right: t=12.39), HES (left: t=12.15; right: t=11.94), STG (left: t=14.39; right: t=14.03), CAL (left: t=13.94; right: t=15.82), CUN (left: t=12.69; right: t=13.64), LING (left: t=13.85; right: t=16.41), IOG (left: t=14.17; right: t=15.82), MOG (left: t=13.13; right: t=12.54), and SOG (left: t=12.90; right: t=11). Subcortical regions displaying high functional connectivity to Pul include AMY (left: t=13.50; right: t=11.75), CAU (left: t=15.24; right: t=16.72), extAMY (left: t=13.41; right: t=13.1), HIP (left: t=11.18; right: t=18.04), HypTH (left: t=11.18; right: t=10.17), MB (left: t=12.14; right: t=10.11), PUT (left: t=13.44; right: t=13.85), SNr (left: t=12.14; right: t=10.17) and left STN (t=11.3). High functional connectivity values were also found for nearly all thalamic nuclei: A (left: t=17.52; right: t=14.83), CM/Pf (left: t=18.66; right: t=15.64), Hb (left: t=18.01; right: t=13.77), MD/CL (left: t=23.6; right: t=18.81), MD (left: t=22.06; right: t=26.03), Ml (left: t=12.9; right: t=16.45), VA (left: t=18.99; right: t=15.67), VL (left: t=25.4; right: t=19.89) and VP (left: t=18.27; right: t=12.76). In addition, high functional connectivity to bilateral cerebellar cortex was also found: CERant (left: 10.84; right: 10.56), CERpost (left: 13.81; right: 14.15), CERfln (t=14.15) and CERpv (t=14.15).

#### Ventral anterior nuclei (VA)

The VA nuclei showed high structural connectivity to right ACCsup (F=1.13%), right MCC (F=1.16%), bilateral IFGpTr (left: F=1.92%; right: F=1.15%), bilateral MFG (left: F=4.06%; right: F=3.67%), bilateral SFG (left: F=7.53%, right: F=6.7%), bilateral dmPFC (left: F=3.07%; right: F=3.99%), and bilateral SMA (left: F=5.04%; right: F=5.88%). Subcortical structures showing high structural connectivity to VA include CAU (left: F=3.26%; right: F=6.54%), left PUT (F=1.69%), RN (left: F=1.61%; right: F=1.29%) and, among brainstem white matter tracts, SCPcr (left: F=2.91%; right: F=2.81%), SCPct (left: 4.64%; right: F=3.84%) and right STT (F=1.16%).

Cortical regions with high functional connectivity to VA include ACCsup (left: t=10.21; right: t=9.31), MCC (left: t=10.93; right: t=10.77), INS (left: t=11.94; right: t=12.54), right IFGpOP (t=10.61), right IFGpOrb (t=10.76), left dmPFC (t=10.93), SMA (left: t=10.6; right: t=10.28), HES (left: t=11.75; right: t=10.32), right TPOsup (t=10.61) and STG (left: t=11.94; right: t=10.87). Among subcortical structures, high peak-t values were found for AMY (left: 10.68; right: t=10.5), CAU (left: t=10.99; right: t=13.54), extAMY (left: t=10.6; right: t=11.24), right HIP (t=10.44), right HypTH (15.26), PUT (left: t=11.57; right: t=11.42), left RN (t=16.82); and STN (left: t=14.89; right: t=11.29). Thalamic nuclei showing high functional connectivity to VA were A (left: 10.23; right: t=16.27), CM/Pf (left: 15.73; right: t=10.16), Hb (left: t=11.68; right: t=10.65), MD/CL (left: t=17.89; right: t=12.04), MD (left: t=14.16, right: t=13.08), left MG (t=10.51), left Ml (t=10.45), Pul (left: t=17.31; right: t=11.56), VL (left: t=19.63; right: t=16.32) and left VP (t=19.63).

#### Ventral lateral nuclei (VL)

High structural connectivity to VL nuclei were found for left IFGpTr (F=1.22%), MFG (left: F=2.71%; right: F=2.48%), SFG (left: F=5.36%; right: F=5.67%), dmPFC (left: F=1.71%; right: F=1.91%), PreCG (left: 3.2%; right: F=5.90%), SMA (left: F=4.88%; right: F=6.62%), PCL (left: F=3.82%; right: F=1.71%) and PoCG (left: F=3%, right: F=2.4%). In addition, VL show high structural connectivity to CAU (left: F=3.05%; right: F=5.48%), PUT (left: F=1.19%), RN (left: F=0.96%; right: F=1.29%), SCPcr (left: F=1.97%; right: F=1.82%), SCPct (left: F=2.81%; right: F=2.67%) and right STT (F=1.23%).

The VL nuclei show high functional connectivity to cortical regions, including ACCpre (left: t=13.35; right: t=11.43), ACCsup (left: t=13.15; right: t=13.35), right MCC (t=16.87), PCC (left: t=17.01; right: t=16.46), INS (left: t=10.13; right: t=11.63), lef OLF (t=12.35), dmPFC (left: t=14.70; right: t=10.11), SMA (left: t=11.45; right: t=9.71), IPL (left: t=10.95; right: t=9.61), left SPL (t=13.54), PCUN (left: t=14.32; right: t=13.62), STG (left: t=11.48; right: t=9.79), CAL (left: t=14.50; right: t=16.08), CUN (left: t=13.27; right: t=13.62), LING (left: t=12.85; right: t=16.08), IOG (left: t=14.49; right:t=16.08), MOG (left: t=13.96; right: t=13.99) and SOG (left: t=14.32; right: t=11.43). Subcortical structures showing high functional connectivity to VL include right AMY (t=12.89), CAU (left: t=22.83; right: t=20.35), extAMY (left: t=17.34; right: t=15.27), HIP (left: t=12.12; right: t=10.66), HypTH (left: t=12.12; right: t=10.24), MB (left: t=11.74; right: t=11.42), PUT (left: t=15.22; right: t=16.59), and left SNr (t=11.74). Among thalamic nuclei, VL obtained high functional connectivity values to left (t=25.05) and right (t=17.57) A, right CM/Pf (t=12.40), left (t=16.49) and right (t=14.66) Hb, left (t=20.25) and right (t=15.77) LD, left (t=15.27) and right (t=18.33) MD/CL, left (t=21.58) and right (t=20.55) MD, left (t=20.52) and right (t=17.01) Ml, left (t=15.23) and right (t=19.50) Pul, and left (t=22.92) and right (t=19.52) VA. VL showed also high functional connectivity to the cerebellar cortex: CERant (left: t=11.78; right: t=11.93), CERpost (left: t=14.5; right: t=14.82), CERfln (t=14.31) and CERpv (t=15.34).

#### Ventral posterior nuclei (VP)

The VP nuclei obtained high structural connectivity values to PreCG (left: F=2.10%; right: F=5.04%), right SMA (F=1.48%), PCL (left: F=4.99%; right: F=2.70%), PoCG (left: F=7.58%; right: F=5.91%), right CAU (F=1.55%), RN (left: F=1.01%; right: F=1.83%), LatL (left: F=1.13%; right: F=2.74%), MedL (left: F=3.5%; right: F=4.02%), SCPcr (left: F=3.53%; right: F=2.95%), SCPct (left: F=4.7%; right: F=4.45%), STT (left: F=3.07%; right: F=4.01), CERant (left: F=2.38%; right: F=2.82%), and CERpost (left: F=1.38%; right: F=1.61%).

High functional connectivity was obtained to MCC (left: t=12.79; right: t=13.16), INS (left: t=16.22; right: t=16.79), IFGpOp (left: t=16.1; right: t=12.88), left IFGpOrb (t=10.95) and left IFGpTr t=11.23), PreCG (left: t=17.11; right: t=18.68), SMA (left: t=14.49; right: t=14.02), PCL (left: t=16.74; right: t=16.42), IPL (left: t=15.19; right: t=12.75), SPL (left: t=12.41; right: t=11.78), PoCG (left: t=18.76; right: t=18.68), PCUN (left: t=12.73; right: t=12.15), ROL (left: t=16.82; right: t=17.13), SMG (left: t=16.79; right: t=17.24); HES (left: t=14.80; right: t=15.05), TPOsup (left: t=14; right: t=14.11), STG (left: t=16.18; right: t=16.53), CAL (left: t=11.21; right: t=11.78), CUN (left: t=13.34; right: t=12.85), LING (left: t=10.96, right: t=11.13) and SOG (left: t=13.33; right: t=12.85). Among thalamic nuclei, high functional connectivity was found to CM/Pf (left: t=22.19; right: t=19.01), MD/CL (left: t=21.18; right: t=14.11), left MD (t=19.66), MG (left: t=12.51; right: t=19.18), Pul (left: t=17.84; right: t=19.44), and VL (left: t=19.45; right: t=19.36). In addition, VP showed high functional connectivity to CERant (left: t=10.27; right: t=10.27).

### Supplementary Figures and Legends

**Supplementary Figure 1.**
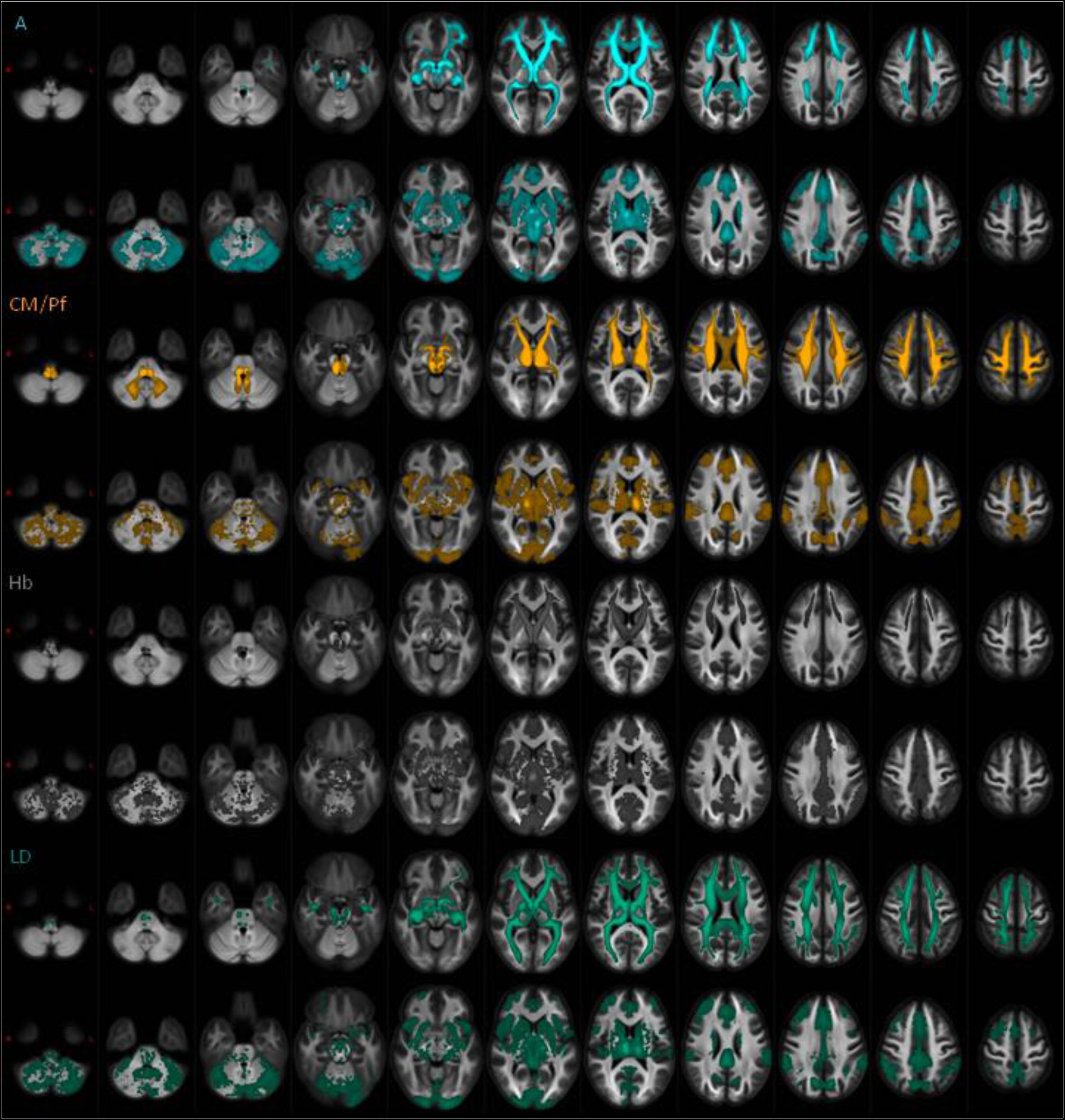
Structural and functional connectivity profiles of thalamic nuclei (A, Hb, CM/Pf, LD). Structural (first row of each nucleus) and functional (second row of each nucleus) group connectivity maps are overlaid on axial sections of the FOD population template (slice increment: 9mm). For visualization purposes, tract MPMs were obtained for each nucleus after weighting of each tractogram according to the sum of streamline weights *f_s_*), application of a 10%, binarization and normalization to FOD template; a 50% threshold was applied to MPMs to show only voxels overlapping in at least half of the sample. Functional activation maps for each subject underwent a random-effect analysis with one-sample t-test, and were masked with a TFCE-corrected p-value < 0.001, then transformed to FOD template space. Details about these processes can be found in text.

**Supplementary Figure 2.**
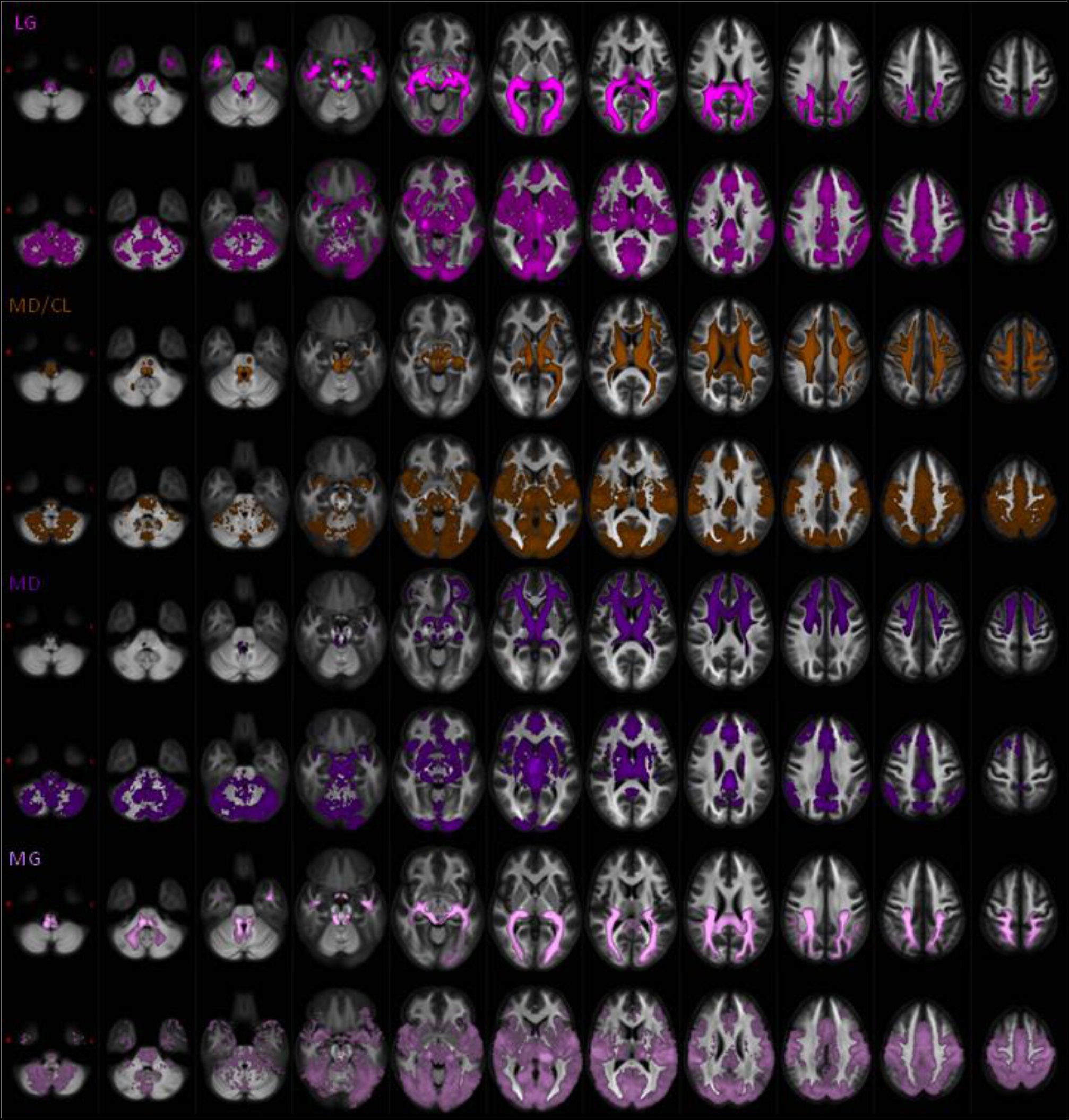
Structural and functional connectivity profiles of thalamic nuclei (LG, MD/CL, MD, MG). Structural (first row of each nucleus) and functional (second row of each nucleus) group connectivity maps are overlaid on axial sections of the FOD population template (slice increment: 9mm). For visualization purposes, tract MPMs were obtained for each nucleus after weighting of each tractogram according to the sum of streamline weights (*f_s_*), application of a 10%, binarization and normalization to FOD template; a 50% threshold was applied to MPMs to show only voxels overlapping in at least half of the sample. Functional activation maps for each subject underwent a random-effect analysis with one-sample t-test, and were masked with a TFCE-corrected p-value < 0.001, then transformed to FOD template space. Details about these processes can be found in text.

**Supplementary Figure 3.**
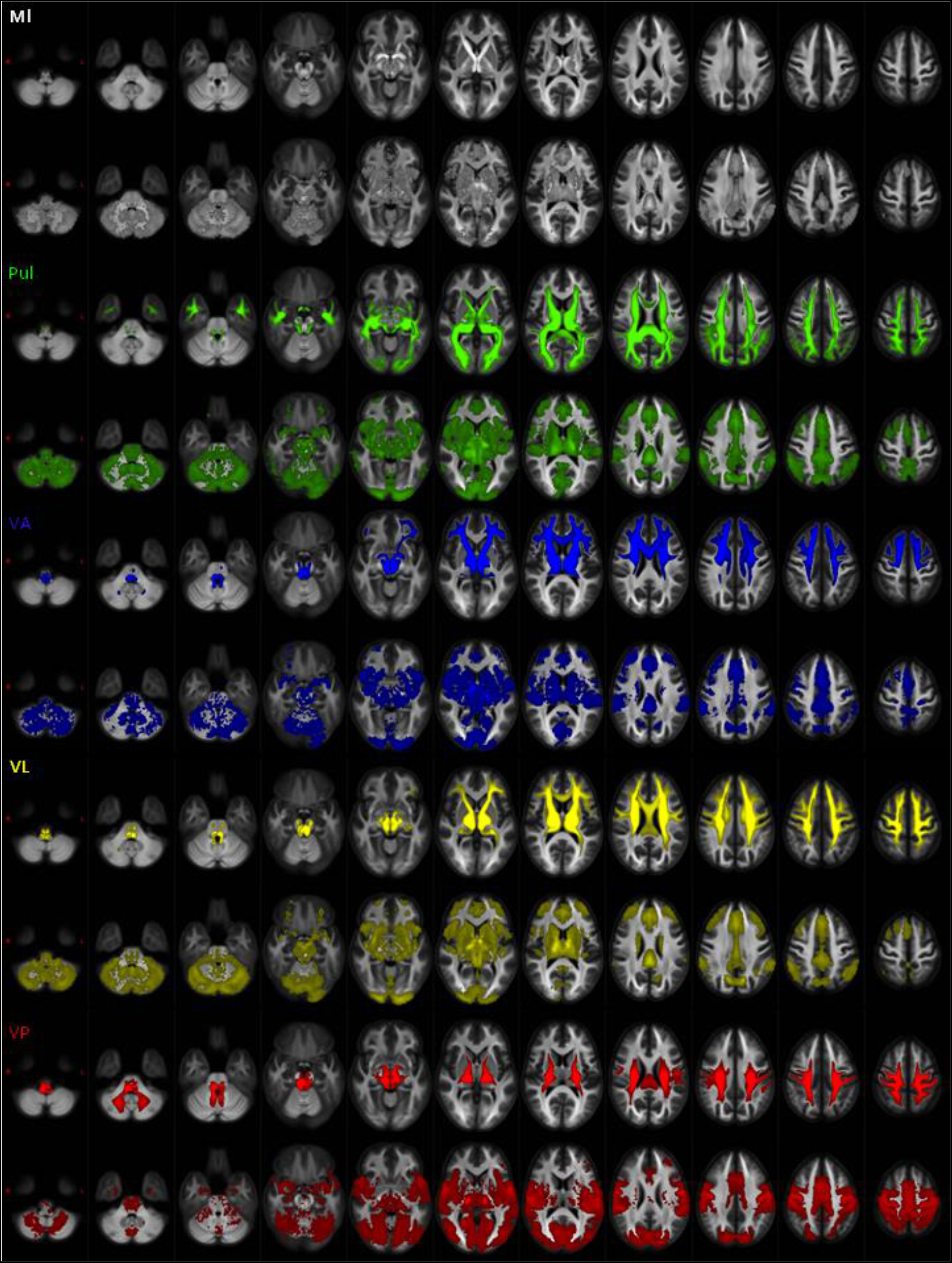
Structural and functional connectivity profiles of thalamic nuclei (Ml, Pul, VA, VL, VP). Structural (first row of each nucleus) and functional (second row of each nucleus) group connectivity maps are overlaid on axial sections of the FOD population template (slice increment: 9mm). For visualization purposes, tract MPMs were obtained for each nucleus after weighting of each tractogram according to the sum of streamline weights (*f_s_*), application of a 10%, binarization and normalization to FOD template; a 50% threshold was applied to MPMs to show only voxels overlapping in at least half of the sample. Functional activation maps for each subject underwent a random-effect analysis with one-sample t-test, and were masked with a TFCE-corrected p-value < 0.001, then transformed to FOD template space. Details about these processes can be found in text.

## Notes

### Competing Interest Statement

The authors have declared no competing interest.

## References

Ackermans L, Duits A, van der Linden C, Tijssen M, Schruers K, Temel Y, Kleijer M, Nederveen P, Bruggeman R, Tromp S, van Kranen-Mastenbroek V, Kingma H, Cath D, Visser-Vandewalle V. 2011. Double-blind clinical trial of thalamic stimulation in patients with Tourette syndrome. Brain. 134:832–844.

Akram H, Dayal V, Mahlknecht P, Georgiev D, Hyam J, Foltynie T, Limousin P, De Vita E, Jahanshahi M, Ashburner J, Behrens T, Hariz M, Zrinzo L. 2018. Connectivity derived thalamic segmentation in deep brain stimulation for tremor. NeuroImage Clin. 18:130–142.

Alexander GE, DeLong MR, Strick PL. 1986. Parallel organization of functionally segregated circuits linking basal ganglia and cortex. Annu Rev Neurosci.

Amin FM, Hougaard A, Magon S, Sprenger T, Wolfram F, Rostrup E, Ashina M. 2018. Altered thalamic connectivity during spontaneous attacks of migraine without aura: A resting-state fMRI study. Cephalalgia. 38:1237–1244.

Andreasen NC. 1997. The Role of the Thalamus in Schizophrenia. Can J Psychiatry. 42:27–33.

Anticevic A, Yang G, Savic A, Murray JD, Cole MW, Repovs G, Pearlson GD, Glahn DC. 2014. Mediodorsal and Visual Thalamic Connectivity Differ in Schizophrenia and Bipolar Disorder With and Without Psychosis History. Schizophr Bull. 40:1227–1243.

Avants BB, Epstein CL, Grossman M, Gee JC. 2008. Symmetric diffeomorphic image registration with cross-correlation: Evaluating automated labeling of elderly and neurodegenerative brain. Med Image Anal.

Balak N, Balkuv E, Karadag A, Basaran R, Biceroglu H, Erkan B, Tanriover N. 2018. Mammillothalamic and Mammillotegmental Tracts as New Targets for Dementia and Epilepsy Treatment. World Neurosurg. 110:133–144.

Banks SD, Coronado RA, Clemons LR, Abraham CM, Pruthi S, Conrad BN, Morgan VL, Guillamondegui OD, Archer KR. 2016. Thalamic Functional Connectivity in Mild Traumatic Brain Injury: Longitudinal Associations With Patient-Reported Outcomes and Neuropsychological Tests. Arch Phys Med Rehabil. 97:1254–1261.

Battistella G, Najdenovska E, Maeder P, Ghazaleh N, Daducci A, Thiran J-P, Jacquemont S, Tuleasca C, Levivier M, Bach Cuadra M, Fornari E. 2017. Robust thalamic nuclei segmentation method based on local diffusion magnetic resonance properties. Brain Struct Funct. 222:2203–2216.

Behrens TEJ, Johansen-Berg H, Woolrich MW, Smith SM, Wheeler-Kingshott CAM, Boulby PA, Barker GJ, Sillery EL, Sheehan K, Ciccarelli O, Thompson AJ, Brady JM, Matthews PM. 2003. Non-invasive mapping of connections between human thalamus and cortex using diffusion imaging. Nat Neurosci. 6:750–757.

Benarroch EE. 2015. Pulvinar: Associative role in cortical function and clinical correlations. Neurology. 84:738–747.

Bertino S, Basile GA, Bramanti A, Anastasi GP, Quartarone A, Milardi D, Cacciola A. 2020. Spatially coherent and topographically organized pathways of the human globus pallidus. Hum Brain Mapp. 41:4641–4661.

Bezdudnaya T, Keller A. 2008. Laterodorsal nucleus of the thalamus: A processor of somatosensory inputs. J Comp Neurol. 507:1979–1989.

Braak H, Braak E. 1991. Alzheimer’s disease affects limbic nuclei of the thalamus. Acta Neuropathol. 81:261–268.

Buckner RL, Krienen FM, Yeo BTT. 2013. Opportunities and limitations of intrinsic functional connectivity MRI. Nat Neurosci. 16:832–837.

Cacciola A, Bertino S, Basile GA, Di Mauro D, Calamuneri A, Chillemi G, Duca A, Bruschetta D, Flace P, Favaloro A, Calabrò RS, Anastasi G, Milardi D. 2019. Mapping the structural connectivity between the periaqueductal gray and the cerebellum in humans. Brain Struct Funct. 224:2153–2165.

Cacciola A, Milardi D, Bertino S, Basile GA, Calamuneri A, Chillemi G, Rizzo G, Anastasi G, Quartarone A. 2019. Structural connectivity-based topography of the human globus pallidus: Implications for therapeutic targeting in movement disorders. Mov Disord. 34:987–996.

Cai S, Huang L, Zou J, Jing L, Zhai B, Ji G, von Deneen KM, Ren J, Ren A. 2015. Changes in Thalamic Connectivity in the Early and Late Stages of Amnestic Mild Cognitive Impairment: A Resting-State Functional Magnetic Resonance Study from ADNI. PLoS One. 10:e0115573.

Calamante F. 2017. Track-weighted imaging methods: extracting information from a streamlines tractogram. Magn Reson Mater Physics, Biol Med. 30:317–335.

Calamante F, Oh S-H, Tournier J-D, Park S-Y, Son Y-D, Chung J-Y, Chi J-G, Jackson GD, Park C-W, Kim Y-B, Connelly A, Cho Z-H. 2013. Super-resolution track-density imaging of thalamic substructures: Comparison with high-resolution anatomical magnetic resonance imaging at 7.0T. Hum Brain Mapp. 34:2538–2548.

Calamante F, Tournier J-D, Heidemann RM, Anwander A, Jackson GD, Connelly A. 2011. Track density imaging (TDI): Validation of super resolution property. Neuroimage. 56:1259–1266.

Calamante F, Tournier J-D, Kurniawan ND, Yang Z, Gyengesi E, Galloway GJ, Reutens DC, Connelly A. 2012. Super-resolution track-density imaging studies of mouse brain: Comparison to histology. Neuroimage. 59:286–296.

Calamante F, Tournier J-D, Smith RE, Connelly A. 2012a. A generalised framework for super-resolution track-weighted imaging. Neuroimage. 59:2494–2503.

Calamante F, Tournier JD, Jackson GD, Connelly A. 2010. Track-density imaging (TDI): Super-resolution white matter imaging using whole-brain track-density mapping. Neuroimage.

Calamante F, Tournier JD, Smith RE, Connelly A. 2012b. A generalised framework for super-resolution track-weighted imaging. Neuroimage.

Chakravarty MM, Bertrand G, Hodge CP, Sadikot AF, Collins DL. 2006. The creation of a brain atlas for image guided neurosurgery using serial histological data. Neuroimage. 30:359–376.

Chakravarty MM, Steadman P, van Eede MC, Calcott RD, Gu V, Shaw P, Raznahan A, Collins DL, Lerch JP. 2013. Performing label-fusion-based segmentation using multiple automatically generated templates. Hum Brain Mapp. 34:2635–2654.

Child ND, Benarroch EE. 2013. Anterior nucleus of the thalamus: Functional organization and clinical implications. Neurology. 81:1869–1876.

Cury RG, Fraix V, Castrioto A, Pérez Fernández MA, Krack P, Chabardes S, Seigneuret E, Alho EJL, Benabid A-L, Moro E. 2017. Thalamic deep brain stimulation for tremor in Parkinson disease, essential tremor, and dystonia. Neurology. 89:1416–1423.

d’Ambrosio A, Hidalgo de la Cruz M, Valsasina P, Pagani E, Colombo B, Rodegher M, Comi G, Filippi M, Rocca MA. 2017. Structural connectivity-defined thalamic subregions have different functional connectivity abnormalities in multiple sclerosis patients: Implications for clinical correlations. Hum Brain Mapp. 38:6005–6018.

da Silva NM, Ahmadi SA, Tafula SN, Cunha JPS, Bötzel K, Vollmar C, Rozanski VE. 2017. A diffusion-based connectivity map of the GPi for optimised stereotactic targeting in DBS. Neuroimage.

Dai J-K, Wang S-X, Shan D, Niu H-C, Lei H. 2018. Super-Resolution Track-Density Imaging Reveals Fine Anatomical Features in Tree Shrew Primary Visual Cortex and Hippocampus. Neurosci Bull. 34:438–448.

DeLong M, Wichmann T. 2010. Changing Views of Basal Ganglia Circuits and Circuit Disorders. Clin EEG Neurosci. 41:61–67.

Deoni SCL, Josseau MJC, Rutt BK, Peters TM. 2005. Visualization of thalamic nuclei on high resolution, multi-averaged T1 and T2 maps acquired at 1.5 T. Hum Brain Mapp. 25:353–359.

Deoni SCL, Rutt BK, Parrent AG, Peters TM. 2007. Segmentation of thalamic nuclei using a modified k-means clustering algorithm and high-resolution quantitative magnetic resonance imaging at 1.5 T. Neuroimage.

Derrington A. 2001. The lateral geniculate nucleus. Curr Biol. 11:R635–R637.

Dhollander T, Emsell L, Van Hecke W, Maes F, Sunaert S, Suetens P. 2014. Track Orientation Density Imaging (TODI) and Track Orientation Distribution (TOD) based tractography. Neuroimage. 94:312–336.

Dhollander T, Raffelt D, Connelly A. 2016. Unsupervised 3-tissue response function estimation from single-shell or multi-shell diffusion MR data without a co-registered T1 image. ISMRM Work Break Barriers Diffus MRI.

Diedrichsen J. 2006. A spatially unbiased atlas template of the human cerebellum. Neuroimage. 33:127–138.

Diedrichsen J, Maderwald S, Küper M, Thürling M, Rabe K, Gizewski ER, Ladd ME, Timmann D. 2011. Imaging the deep cerebellar nuclei: A probabilistic atlas and normalization procedure. Neuroimage. 54:1786–1794.

Eickhoff SB, Thirion B, Varoquaux G, Bzdok D. 2015. Connectivity-based parcellation: Critique and implications. Hum Brain Mapp.

Erica T, White M, Zrinzo L. 2014. Abstracts. Clin Anat. 27:262–273.

Ewert S, Plettig P, Li N, Chakravarty MM, Collins DL, Herrington TM, Kühn AA, Horn A. 2018. Toward defining deep brain stimulation targets in MNI space: A subcortical atlas based on multimodal MRI, histology and structural connectivity. Neuroimage. 170:271–282.

Fan Y, Nickerson LD, Li H, Ma Y, Lyu B, Miao X, Zhuo Y, Ge J, Zou Q, Gao J-H. 2015. Functional Connectivity-Based Parcellation of the Thalamus: An Unsupervised Clustering Method and Its Validity Investigation. Brain Connect. 5:620–630.

Fang P-C, Stepniewska I, Kaas JH. 2006. The thalamic connections of motor, premotor, and prefrontal areas of cortex in a prosimian primate (Otolemur garnetti). Neuroscience. 143:987–1020.

Fiechter M, Nowacki A, Oertel MF, Fichtner J, Debove I, Lachenmayer ML, Wiest R, Bassetti CL, Raabe A, Kaelin-Lang A, Schüpbach MW, Pollo C. 2017. Deep Brain Stimulation for Tremor: Is There a Common Structure? Stereotact Funct Neurosurg. 95:243–250.

Fisher R, Salanova V, Witt T, Worth R, Henry T, Gross R, Oommen K, Osorio I, Nazzaro J, Labar D, Kaplitt M, Sperling M, Sandok E, Neal J, Handforth A, Stern J, DeSalles A, Chung S, Shetter A, Bergen D, Bakay R, Henderson J, French J, Baltuch G, Rosenfeld W, Youkilis A, Marks W, Garcia P, Barbaro N, Fountain N, Bazil C, Goodman R, McKhann G, Babu Krishnamurthy K, Papavassiliou S, Epstein C, Pollard J, Tonder L, Grebin J, Coffey R, Graves N. 2010. Electrical stimulation of the anterior nucleus of thalamus for treatment of refractory epilepsy. Epilepsia. 51:899–908.

Fonov V, Evans A, McKinstry R, Almli C, Collins D. 2009. Unbiased nonlinear average age-appropriate brain templates from birth to adulthood. Neuroimage.

Glasser MF, Sotiropoulos SN, Wilson JA, Coalson TS, Fischl B, Andersson JL, Xu J, Jbabdi S, Webster M, Polimeni JR, Van Essen DC, Jenkinson M. 2013. The minimal preprocessing pipelines for the Human Connectome Project. Neuroimage. 80:105–124.

Granziera C, Daducci A, Romascano D, Roche A, Helms G, Krueger G, Hadjikhani N. 2014. Structural abnormalities in the thalamus of migraineurs with aura: A multiparametric study at 3 T. Hum Brain Mapp. 35:1461–1468.

Groenewegen HJ, Berendse HW, Haber SN. 1993. Organization of the output of the ventral striatopallidal system in the rat: Ventral pallidal efferents. Neuroscience. 57:113–142.

Hassler. 1983. Stereotaxy of the human brain — anatomical, physiological and clinical applications. Clin Neurol Neurosurg.

Hassler R. 1959. Anatomy of the Thalamus. In: Schaltenbrand G,, Bayley P, editors. Introduction to Stereotaxic Operations With an Atlas of the Human Brain. 1st ed. Stuttgart; New York, NY: Georg Thieme Verlag. p. 230–290.

Henderson JM. 2000. Loss of thalamic intralaminar nuclei in progressive supranuclear palsy and Parkinson’s disease: clinical and therapeutic implications. Brain. 123:1410–1421.

Herkenham M, Nauta WJH. 1979. Efferent connections of the habenular nuclei in the rat. J Comp Neurol. 187:19–47.

Hirai T, Jones EG. 1989. A new parcellation of the human thalamus on the basis of histochemical staining. Brain Res Rev. 14:1–34.

Hongzhi Wang, Suh JW, Das SR, Pluta JB, Craige C, Yushkevich PA. 2013. Multi-Atlas Segmentation with Joint Label Fusion. IEEE Trans Pattern Anal Mach Intell. 35:611–623.

Hwang K, Bertolero MA, Liu WB, D’Esposito M. 2017. The Human Thalamus Is an Integrative Hub for Functional Brain Networks. J Neurosci. 37:5594–5607.

Iglehart C, Monti M, Cain J, Tourdias T, Saranathan M. 2020. A systematic comparison of structural-, structural connectivity-, and functional connectivity-based thalamus parcellation techniques. Brain Struct Funct. 225:1631–1642.

Iglesias JE, Insausti R, Lerma-Usabiaga G, Bocchetta M, Van Leemput K, Greve DN, van der Kouwe A, Fischl B, Caballero-Gaudes C, Paz-Alonso PM. 2018a. A probabilistic atlas of the human thalamic nuclei combining ex vivo MRI and histology. Neuroimage.

Iglesias JE, Insausti R, Lerma-Usabiaga G, Bocchetta M, Van Leemput K, Greve DN, van der Kouwe A, Fischl B, Caballero-Gaudes C, Paz-Alonso PM. 2018b. A probabilistic atlas of the human thalamic nuclei combining ex vivo MRI and histology. Neuroimage. 183:314–326.

Ilinsky I, Horn A, Paul-Gilloteaux P, Gressens P, Verney C, Kultas-Ilinsky K. 2018. Human Motor Thalamus Reconstructed in 3D from Continuous Sagittal Sections with Identified Subcortical Afferent Territories. eneuro. 5:ENEURO.0060-18.2018.

Jain AK. 2010. Data clustering: 50 years beyond K-means. Pattern Recognit Lett. 31:651–666.

Jeurissen B, Tournier J-D, Dhollander T, Connelly A, Sijbers J. 2014a. Multi-tissue constrained spherical deconvolution for improved analysis of multi-shell diffusion MRI data. Neuroimage. 103:411–426.

Jeurissen B, Tournier JD, Dhollander T, Connelly A, Sijbers J. 2014b. Multi-tissue constrained spherical deconvolution for improved analysis of multi-shell diffusion MRI data. Neuroimage.

Ji B, Li Z, Li K, Li L, Langley J, Shen H, Nie S, Zhang R, Hu X. 2016. Dynamic thalamus parcellation from resting-state fMRI data. Hum Brain Mapp. 37:954–967.

Jiltsova E, Möttönen T, Fahlström M, Haapasalo J, Tähtinen T, Peltola J, Öhman J, Larsson EM, Kiekara T, Lehtimäki K. 2016. Imaging of Anterior Nucleus of Thalamus Using 1.5T MRI for Deep Brain Stimulation Targeting in Refractory Epilepsy. Neuromodulation.

Johansen-Berg H, Behrens TEJ, Sillery E, Ciccarelli O, Thompson AJ, Smith SM, Matthews PM. 2005. Functional–Anatomical Validation and Individual Variation of Diffusion Tractography-based Segmentation of the Human Thalamus. Cereb Cortex. 15:31–39.

Jones DK, Knösche TR, Turner R. 2013. White matter integrity, fiber count, and other fallacies: The do’s and don’ts of diffusion MRI. Neuroimage. 73:239–254.

Kanowski M, Voges J, Buentjen L, Stadler J, Heinze HJ, Tempelmann C. 2014. Direct Visualization of Anatomic Subfields within the Superior Aspect of the Human Lateral Thalamus by MRI at 7T. Am J Neuroradiol. 35:1721–1727.

Krack P, Dostrovsky J, Ilinsky I, Kultas-Ilinsky K, Lenz F, Lozano A, Vitek J. 2002. Surgery of the motor thalamus: Problems with the present nomenclatures. Mov Disord. 17:S2–S8.

Krauth A, Blanc R, Poveda A, Jeanmonod D, Morel A, Székely G. 2010. A mean three-dimensional atlas of the human thalamus: Generation from multiple histological data. Neuroimage.

Kultas-Ilinsky K, Sivan-Loukianova E, Ilinsky IA. 2003. Reevaluation of the primary motor cortex connections with the thalamus in primates. J Comp Neurol. 457:133–158.

Kumar V, Mang S, Grodd W. 2015. Direct diffusion-based parcellation of the human thalamus. Brain Struct Funct. 220:1619–1635.

Kumar VJ, van Oort E, Scheffler K, Beckmann CF, Grodd W. 2017. Functional anatomy of the human thalamus at rest. Neuroimage. 147:678–691.

Lambert C, Simon H, Colman J, Barrick TR. 2017. Defining thalamic nuclei and topographic connectivity gradients in vivo. Neuroimage. 158:466–479.

Leh SE, Chakravarty MM, Ptito A. 2008. The Connectivity of the Human Pulvinar: A Diffusion Tensor Imaging Tractography Study. Int J Biomed Imaging. 2008:1–5.

Lin F, Zivadinov R, Hagemeier J, Weinstock-Guttman B, Vaughn C, Gandhi S, Jakimovski D, Hulst HE, Benedict RHB, Bergsland N, Fuchs T, Dwyer MG. 2019. Altered nuclei-specific thalamic functional connectivity patterns in multiple sclerosis and their associations with fatigue and cognition. Mult Scler J. 25:1243–1254.

Liu Y, D’Haese P-F, Newton AT, Dawant BM. 2020. Generation of human thalamus atlases from 7 T data and application to intrathalamic nuclei segmentation in clinical 3 T T1-weighted images. Magn Reson Imaging. 65:114–128.

Maffei C, Sarubbo S, Jovicich J. 2019. Diffusion-based tractography atlas of the human acoustic radiation. Sci Rep.

Mai JK, Forutan F. 2012. Thalamus. In: The Human Nervous System. Elsevier. p. 618–677.

Mai JK, Majtanik M. 2019. Toward a Common Terminology for the Thalamus. Front Neuroanat. 12.

Maier-Hein KH, Neher PF, Houde J-C, Côté M-A, Garyfallidis E, Zhong J, Chamberland M, Yeh F-C, Lin Y-C, Ji Q, Reddick WE, Glass JO, Chen DQ, Feng Y, Gao C, Wu Y, Ma J, He R, Li Q, Westin C-F, Deslauriers-Gauthier S, González JOO, Paquette M, St-Jean S, Girard G, Rheault F, Sidhu J, Tax CMW, Guo F, Mesri HY, Dávid S, Froeling M, Heemskerk AM, Leemans A, Boré A, Pinsard B, Bedetti C, Desrosiers M, Brambati S, Doyon J, Sarica A, Vasta R, Cerasa A, Quattrone A, Yeatman J, Khan AR, Hodges W, Alexander S, Romascano D, Barakovic M, Auría A, Esteban O, Lemkaddem A, Thiran J-P, Cetingul HE, Odry BL, Mailhe B, Nadar MS, Pizzagalli F, Prasad G, Villalon-Reina JE, Galvis J, Thompson PM, Requejo FDS, Laguna PL, Lacerda LM, Barrett R, Dell’Acqua F, Catani M, Petit L, Caruyer E, Daducci A, Dyrby TB, Holland-Letz T, Hilgetag CC, Stieltjes B, Descoteaux M. 2017. The challenge of mapping the human connectome based on diffusion tractography. Nat Commun. 8:1349.

McFarland NR, Haber SN. 2002. Thalamic Relay Nuclei of the Basal Ganglia Form Both Reciprocal and Nonreciprocal Cortical Connections, Linking Multiple Frontal Cortical Areas. J Neurosci. 22:8117–8132.

Middlebrooks EH, Tuna IS, Almeida L, Grewal SS, Wong J, Heckman MG, Lesser ER, Bredel M, Foote KD, Okun MS, Holanda VM. 2018. Structural connectivity–based segmentation of the thalamus and prediction of tremor improvement following thalamic deep brain stimulation of the ventral intermediate nucleus. NeuroImage Clin. 20:1266–1273.

Milardi D, Quartarone A, Bramanti A, Anastasi G, Bertino S, Basile GA, Buonasera P, Pilone G, Celeste G, Rizzo G, Bruschetta D, Cacciola A. 2019. The Cortico-Basal Ganglia-Cerebellar Network: Past, Present and Future Perspectives. Front Syst Neurosci.

Milosevic L, Kalia SK, Hodaie M, Lozano AM, Popovic MR, Hutchison WD. 2018. Physiological mechanisms of thalamic ventral intermediate nucleus stimulation for tremor suppression. Brain. 141:2142–2155.

Mitchell AS, Chakraborty S. 2013. What does the mediodorsal thalamus do? Front Syst Neurosci. 7.

Mitchell AS, Dalrymple-Alford JC. 2005. Dissociable memory effects after medial thalamus lesions in the rat. Eur J Neurosci. 22:973–985.

Mizumori SJY, Williams JD. 1993. Directionally selective mnemonic properties of neurons in the lateral dorsal nucleus of the thalamus of rats. J Neurosci.

Morel A, Magnin M, Jeanmonod D. 1997. Multiarchitectonic and stereotactic atlas of the human thalamus. J Comp Neurol. 387:588–630.

Möttönen T, Katisko J, Haapasalo J, Tähtinen T, Kiekara T, Kähärä V, Peltola J, Öhman J, Lehtimäki K. 2015. Defining the anterior nucleus of the thalamus (ANT) as a deep brain stimulation target in refractory epilepsy: Delineation using 3 T MRI and intraoperative microelectrode recording. NeuroImage Clin. 7:823–829.

Murray JD, Anticevic A. 2017. Toward understanding thalamocortical dysfunction in schizophrenia through computational models of neural circuit dynamics. Schizophr Res.

Najdenovska E, Alemán-Gómez Y, Battistella G, Descoteaux M, Hagmann P, Jacquemont S, Maeder P, Thiran J-P, Fornari E, Bach Cuadra M. 2018. In-vivo probabilistic atlas of human thalamic nuclei based on diffusion-weighted magnetic resonance imaging. Sci Data. 5:180270.

O’Muircheartaigh J, Vollmar C, Traynor C, Barker GJ, Kumari V, Symms MR, Thompson P, Duncan JS, Koepp MJ, Richardson MP. 2011. Clustering probabilistic tractograms using independent component analysis applied to the thalamus. Neuroimage. 54:2020–2032.

Ossowska K. 2020. Zona incerta as a therapeutic target in Parkinson’s disease. J Neurol. 267:591–606.

Owens-Walton C, Jakabek D, Power BD, Walterfang M, Velakoulis D, van Westen D, Looi JCL, Shaw M, Hansson O. 2019. Increased functional connectivity of thalamic subdivisions in patients with Parkinson’s disease. PLoS One. 14:e0222002.

Padberg J, Cerkevich C, Engle J, Rajan AT, Recanzone G, Kaas J, Krubitzer L. 2009. Thalamocortical Connections of Parietal Somatosensory Cortical Fields in Macaque Monkeys are Highly Divergent and Convergent. Cereb Cortex. 19:2038–2064.

Paez P, Crow T, Mackay C. 2011. Fronto-temporal connections in schizophrenia and bipolar disorder. Int Clin Psychopharmacol. 26:e143.

Parkes L, Fulcher B, Yücel M, Fornito A. 2018. An evaluation of the efficacy, reliability, and sensitivity of motion correction strategies for resting-state functional MRI. Neuroimage. 171:415–436.

Parnaudeau S, Bolkan SS, Kellendonk C. 2018. The Mediodorsal Thalamus: An Essential Partner of the Prefrontal Cortex for Cognition. Biol Psychiatry. 83:648–656.

Pauli WM, Nili AN, Tyszka JM. 2018. A high-resolution probabilistic in vivo atlas of human subcortical brain nuclei. Sci Data. 5:180063.

Percheron G. 2003. Thalamus. In: The Human Nervous System: Second Edition.

Percheron G, François C, Talbi B, Yelnik J, Fénelon G. 1996. The primate motor thalamus. Brain Res Rev. 22:93–181.

Pergola G, Selvaggi P, Trizio S, Bertolino A, Blasi G. 2015. The role of the thalamus in schizophrenia from a neuroimaging perspective. Neurosci Biobehav Rev. 54:57–75.

Pietsch M, Raffelt D, Dhollander T, Tournier J-D. 2017. Multi-contrast diffeomorphic non-linear registration of orientation density functions. In: 25th International Society of Magnetic Resonance in Medicine. p. 3522.

Plachti A, Eickhoff SB, Hoffstaedter F, Patil KR, Laird AR, Fox PT, Amunts K, Genon S. 2019. Multimodal Parcellations and Extensive Behavioral Profiling Tackling the Hippocampus Gradient. Cereb Cortex. 29:4595–4612.

Planche V, Su JH, Mournet S, Saranathan M, Dousset V, Han M, Rutt BK, Tourdias T. 2020. White-matter-nulled MPRAGE at 7T reveals thalamic lesions and atrophy of specific thalamic nuclei in multiple sclerosis. Mult Scler J. 26:987–992.

Planetta PJ, Schulze ET, Geary EK, Corcos DM, Goldman JG, Little DM, Vaillancourt DE. 2013. Thalamic Projection Fiber Integrity in de novo Parkinson Disease. Am J Neuroradiol. 34:74–79.

Preller KH, Burt JB, Ji JL, Schleifer CH, Adkinson BD, Stämpfli P, Seifritz E, Repovs G, Krystal JH, Murray JD, Vollenweider FX, Anticevic A. 2018. Changes in global and thalamic brain connectivity in LSD-induced altered states of consciousness are attributable to the 5-HT2A receptor. Elife. 7.

Quina LA, Tempest L, Ng L, Harris JA, Ferguson S, Jhou TC, Turner EE. 2015. Efferent Pathways of the Mouse Lateral Habenula. J Comp Neurol. 523:32–60.

Raffelt D, Tournier J-D, Crozier S, Connelly A, Salvado O. 2012. Reorientation of fiber orientation distributions using apodized point spread functions. Magn Reson Med. 67:844–855.

Raffelt D, Tournier J-D, Fripp J, Crozier S, Connelly A, Salvado O. 2011. Symmetric diffeomorphic registration of fibre orientation distributions. Neuroimage. 56:1171–1180.

Ramsay IS. 2019. An Activation Likelihood Estimate Meta-analysis of Thalamocortical Dysconnectivity in Psychosis. Biol Psychiatry Cogn Neurosci Neuroimaging. 4:859–869.

Rolls ET, Huang C-C, Lin C-P, Feng J, Joliot M. 2020. Automated anatomical labelling atlas 3. Neuroimage. 206:116189.

Rossi M, Cerquetti D, Mandolesi J, Merello M. 2016. Thalamotomy, pallidotomy and subthalamotomy in the management of Parkinson’s disease. In: Galvez-Jimenez N,, Fernandez HH,, Espay AJ,, Fox SH, editors. Parkinson’s Disease. Cambridge: Cambridge University Press. p. 175–186.

Saalmann YB. 2014. Intralaminar and medial thalamic influence on cortical synchrony, information transmission and cognition. Front Syst Neurosci. 8.

Sadikot AF, Chakravarty MM, Bertrand G, Rymar V V., Al-Subaie F, Collins DL. 2011. Creation of Computerized 3D MRI-Integrated Atlases of the Human Basal Ganglia and Thalamus. Front Syst Neurosci. 5.

Salimi-Khorshidi G, Douaud G, Beckmann CF, Glasser MF, Griffanti L, Smith SM. 2014. Automatic denoising of functional MRI data: Combining independent component analysis and hierarchical fusion of classifiers. Neuroimage. 90:449–468.

Shamir RR, Duchin Y, Kim J, Sapiro G, Harel N. 2016. Segmentation overlap measures are biased to structure’s size but correctable. Int J Comput Assist Radiol Surg. S44–S45.

Shelton L, Pendse G, Maleki N, Moulton EA, Lebel A, Becerra L, Borsook D. 2012. Mapping pain activation and connectivity of the human habenula. J Neurophysiol. 107:2633–2648.

Sitek KR, Gulban OF, Calabrese E, Johnson GA, Lage-Castellanos A, Moerel M, Ghosh SS, De Martino F. 2019. Mapping the human subcortical auditory system using histology, postmortem MRI and in vivo MRI at 7T. Elife. 8.

Smith RE, Tournier J-D, Calamante F, Connelly A. 2012. Anatomically-constrained tractography: Improved diffusion MRI streamlines tractography through effective use of anatomical information. Neuroimage. 62:1924–1938.

Smith RE, Tournier J-D, Calamante F, Connelly A. 2015. SIFT2: Enabling dense quantitative assessment of brain white matter connectivity using streamlines tractography. Neuroimage. 119:338–351.

Smith S, Nichols T. 2009. Threshold-free cluster enhancement: Addressing problems of smoothing, threshold dependence and localisation in cluster inference. Neuroimage. 44:83–98.

Smith SM, Beckmann CF, Andersson J, Auerbach EJ, Bijsterbosch J, Douaud G, Duff E, Feinberg DA, Griffanti L, Harms MP, Kelly M, Laumann T, Miller KL, Moeller S, Petersen S, Power J, Salimi-Khorshidi G, Snyder AZ, Vu AT, Woolrich MW, Xu J, Yacoub E, Uğurbil K, Van Essen DC, Glasser MF. 2013. Resting-state fMRI in the Human Connectome Project. Neuroimage. 80:144–168.

Son B, Shon YM, Choi J, Kim J, Ha S, Kim S-H, Lee S. 2016. Clinical Outcome of Patients with Deep Brain Stimulation of the Centromedian Thalamic Nucleus for Refractory Epilepsy and Location of the Active Contacts. Stereotact Funct Neurosurg. 94:187–197.

Sotiropoulos SN, Jbabdi S, Xu J, Andersson JL, Moeller S, Auerbach EJ, Glasser MF, Hernandez M, Sapiro G, Jenkinson M, Feinberg DA, Yacoub E, Lenglet C, Van Essen DC, Ugurbil K, Behrens TEJ, WU-Minn HCP Consortium. 2013. Advances in diffusion MRI acquisition and processing in the Human Connectome Project. Neuroimage. 80:125–143.

Spiegelmann R, Nissim O, Daniels D, Ocherashvilli A, Mardor Y. 2006. Stereotactic Targeting of the Ventrointermediate Nucleus of the Thalamus by Direct Visualization with High-Field MRI. Stereotact Funct Neurosurg. 84:19–23.

Stepniewska I, Preuss TM, Kaas JH. 1994. Architectonic subdivisions of the motor thalamus of owl monkeys: Nissl, acetylcholinesterase, and cytochrome oxidase patterns. J Comp Neurol. 349:536–557.

Stoodley CJ, Schmahmann JD. 2018. Functional topography of the human cerebellum. In: Handbook of Clinical Neurology. p. 59–70.

Strotmann B, Heidemann RM, Anwander A, Weiss M, Trampel R, Villringer A, Turner R. 2014. High-resolution MRI and diffusion-weighted imaging of the human habenula at 7 tesla. J Magn Reson Imaging. 39:1018–1026.

Su JH, Thomas FT, Kasoff WS, Tourdias T, Choi EY, Rutt BK, Saranathan M. 2019. Thalamus Optimized Multi Atlas Segmentation (THOMAS): fast, fully automated segmentation of thalamic nuclei from structural MRI. Neuroimage. 194:272–282.

Sudhyadhom A, Haq IU, Foote KD, Okun MS, Bova FJ. 2009. A high resolution and high contrast MRI for differentiation of subcortical structures for DBS targeting: The Fast Gray Matter Acquisition T1 Inversion Recovery (FGATIR). Neuroimage. 47:T44–T52.

Tang Y, Sun W, Toga AW, Ringman JM, Shi Y. 2018. A probabilistic atlas of human brainstem pathways based on connectome imaging data. Neuroimage. 169:227–239.

Tourdias T, Saranathan M, Levesque IR, Su J, Rutt BK. 2014. Visualization of intra-thalamic nuclei with optimized white-matter-nulled MPRAGE at 7T. Neuroimage. 84:534–545.

Tournier J-D, Calamante F, Connelly A. 2010. Improved probabilistic streamlines tractography by 2nd order integration over fibre orientation distributions. Proc Int Soc Magn Reson Med.

Tournier J-D, Calamante F, Connelly A. 2012. MRtrix: Diffusion tractography in crossing fiber regions. Int J Imaging Syst Technol. 22:53–66.

Tournier J-D, Yeh C-H, Calamante F, Cho K-H, Connelly A, Lin C-P. 2008. Resolving crossing fibres using constrained spherical deconvolution: Validation using diffusion-weighted imaging phantom data. Neuroimage. 42:617–625.

Traynor C, Heckemann RA, Hammers A, O’Muircheartaigh J, Crum WR, Barker GJ, Richardson MP. 2010. Reproducibility of thalamic segmentation based on probabilistic tractography. Neuroimage. 52:69–85.

Traynor CR, Barker GJ, Crum WR, Williams SCR, Richardson MP. 2011. Segmentation of the thalamus in MRI based on T1 and T2. Neuroimage. 56:939–950.

Treiber JM, White NS, Steed TC, Bartsch H, Holland D, Farid N, McDonald CR, Carter BS, Dale AM, Chen CC. 2016. Characterization and Correction of Geometric Distortions in 814 Diffusion Weighted Images. PLoS One. 11:e0152472.

Uğurbil K, Xu J, Auerbach EJ, Moeller S, Vu AT, Duarte-Carvajalino JM, Lenglet C, Wu X, Schmitter S, Van de Moortele PF, Strupp J, Sapiro G, De Martino F, Wang D, Harel N, Garwood M, Chen L, Feinberg DA, Smith SM, Miller KL, Sotiropoulos SN, Jbabdi S, Andersson JLR, Behrens TEJ, Glasser MF, Van Essen DC, Yacoub E. 2013. Pushing spatial and temporal resolution for functional and diffusion MRI in the Human Connectome Project. Neuroimage. 80:80–104.

Van der Werf YD, Witter MP, Groenewegen HJ. 2002. The intralaminar and midline nuclei of the thalamus. Anatomical and functional evidence for participation in processes of arousal and awareness. Brain Res Rev. 39:107–140.

Van Essen DC, Smith SM, Barch DM, Behrens TEJ, Yacoub E, Ugurbil K. 2013. The WU-Minn Human Connectome Project: An overview. Neuroimage.

Van Essen DC, Ugurbil K, Auerbach E, Barch D, Behrens TEJ, Bucholz R, Chang A, Chen L, Corbetta M, Curtiss SW, Della Penna S, Feinberg D, Glasser MF, Harel N, Heath AC, Larson-Prior L, Marcus D, Michalareas G, Moeller S, Oostenveld R, Petersen SE, Prior F, Schlaggar BL, Smith SM, Snyder AZ, Xu J, Yacoub E. 2012a. The Human Connectome Project: A data acquisition perspective. Neuroimage. 62:2222–2231.

Van Essen DC, Ugurbil K, Auerbach E, Barch D, Behrens TEJ, Bucholz R, Chang A, Chen L, Corbetta M, Curtiss SW, Della Penna S, Feinberg D, Glasser MF, Harel N, Heath AC, Larson-Prior L, Marcus D, Michalareas G, Moeller S, Oostenveld R, Petersen SE, Prior F, Schlaggar BL, Smith SM, Snyder AZ, Xu J, Yacoub E. 2012b. The Human Connectome Project: A data acquisition perspective. Neuroimage. 62:2222–2231.

van Groen T, Kadish I, Wyss JM. 2002. The role of the laterodorsal nucleus of the thalamus in spatial learning and memory in the rat. Behav Brain Res. 136:329–337.

van Oort ESB, Mennes M, Navarro Schröder T, Kumar VJ, Zaragoza Jimenez NI, Grodd W, Doeller CF, Beckmann CF. 2018. Functional parcellation using time courses of instantaneous connectivity. Neuroimage. 170:31–40.

Vandewalle V, van der Linden C, Groenewegen H, Caemaert J. 1999. Stereotactic treatment of Gilles de la Tourette syndrome by high frequency stimulation of thalamus. Lancet. 353:724.

Vassal F, Coste J, Derost P, Mendes V, Gabrillargues J, Nuti C, Durif F, Lemaire J-J. 2012. Direct stereotactic targeting of the ventrointermediate nucleus of the thalamus based on anatomic 1.5-T MRI mapping with a white matter attenuated inversion recovery (WAIR) sequence. Brain Stimul. 5:625–633.

Vernaleken I, Kuhn J, Lenartz D, Raptis M, Huff W, Janouschek H, Neuner I, Schaefer WM, Gründer G, Sturm V. 2009. Bithalamical Deep Brain Stimulation in Tourette Syndrome Is Associated with Reduction in Dopaminergic Transmission. Biol Psychiatry. 66:e15–e17.

Vertes RP, Linley SB, Hoover WB. 2015. Limbic circuitry of the midline thalamus. Neurosci Biobehav Rev. 54:89–107.

Wang Z, Jia X, Liang P, Qi Z, Yang Y, Zhou W, Li K. 2012. Changes in thalamus connectivity in mild cognitive impairment: Evidence from resting state fMRI. Eur J Radiol. 81:277–285.

Watanabe Y, Funahashi S. 2012. Thalamic mediodorsal nucleus and working memory. Neurosci Biobehav Rev.

Whitfield-Gabrieli S, Nieto-Castanon A. 2012. Conn: A Functional Connectivity Toolbox for Correlated and Anticorrelated Brain Networks. Brain Connect. 2:125–141.

Winer JA. 1984. The human medial geniculate body. Hear Res. 15:225–247.

Yang S, Meng Y, Li J, Li B, Fan Y-S, Chen H, Liao W. 2020. The thalamic functional gradient and its relationship to structural basis and cognitive relevance. Neuroimage. 218:116960.

Yin Y, Jin C, Hu X, Duan L, Li Z, Song M, Chen H, Feng B, Jiang T, Jin H, Wong C, Gong Q, Li L. 2011. Altered resting-state functional connectivity of thalamus in earthquake-induced posttraumatic stress disorder: A functional magnetic resonance imaging study. Brain Res. 1411:98–107.

Young RF, Jacques DS, Rand RW, Copcutt BR. 1994. Medial Thalamotomy with the Leksell Gamma Knife for Treatment of Chronic Pain. In: Acta neurochirurgica. Supplement. p. 105–110.

Zhang D, Snyder AZ, Fox MD, Sansbury MW, Shimony JS, Raichle ME. 2008. Intrinsic Functional Relations Between Human Cerebral Cortex and Thalamus. J Neurophysiol. 100:1740–1748.

Zhang D, Snyder AZ, Shimony JS, Fox MD, Raichle ME. 2010. Noninvasive Functional and Structural Connectivity Mapping of the Human Thalamocortical System. Cereb Cortex. 20:1187–1194.

Zhou B, Liu Y, Zhang Z, An N, Yao H, Wang P, Wang L, Zhang X, Jiang T. 2013. Impaired Functional Connectivity of the Thalamus in Alzheimer’s Disease and Mild Cognitive Impairment: A Resting-State fMRI Study. Curr Alzheimer Res. 10:754–766.

